# Modular Albumin-Chaperoned NIR-II Nanofluorophores Enables Pan-Ovarian Cancer Imaging Across Multiscale Tumor Models

**DOI:** 10.64898/2026.05.06.717945

**Authors:** Isabella Vasquez, Lucia L. Nash, Md Shahriar, Mahenour Megahed, Jerry Cao-Xue, Itzel De León, Changxue Xu, Ulrich Bickel, Shreya A. Raghavan, Indrajit Srivastava

## Abstract

Ovarian cancer remains the most lethal gynecological malignancy, primarily due to late-stage diagnosis and the challenges of achieving complete cytoreduction. While fluorescence image-guided surgery (FIGS) offers intraoperative visualization, current clinical agents are limited by insufficient brightness, rapid photobleaching, and poor molecular selectivity, particularly in the near-infrared window. Here, we report the rational modular design of ultrabright NIR-II semiconducting polymer (SP) nanofluorophores for high-fidelity ovarian cancer imaging. By nanoconfining of a representative hydrophobic SP within a functional albumin matrix induces a “chaperone” effect that suppresses aggregation-induced quenching and shifts emission in the NIR-II window (1000-1250 nm). This platform integrates a dual-receptor targeting strategy, leveraging intrinsic albumin-receptor interactions (GP60 and SPARC) alongside folate receptor alpha (FRα) functionalization. This synergistic approach enables pan-ovarian cancer imaging by ensuring high-affinity binding across diverse tumor phenotypes, regardless of heterogeneous receptor expression. Across a multiscale validation framework, the nanofluorophores demonstrate efficient receptor-mediated endocytosis in 2D cultures and deep interstitial penetration in 3D tumor spheroids. Furthermore, microfluidic tumor-on-chip models incorporating endothelial-like fenestrations confirm controlled extravasation and targeting under physiological shear stress. 3D bioprinted tumor phantoms and *ex vivo* porcine ovary tissues further confirm that BSA-FA@SP2 provides superior lesion delineation and signal-to-background ratios compared to indocyanine green, a clinical standard. Importantly, the nanofluorophores exhibit excellent hemocompatibility, with minimal hemolysis and negligible complement activation, indicating a non-immunogenic, stealth profile. Collectively, this work establishes albumin-shielded NIR-II nanofluorophores as a robust platform for precision intraoperative pan-ovarian imaging and advances the translational potential of nanotechnology-enabled surgical oncology.

## 1. Introduction

Ovarian cancer remains the deadliest gynecological cancer, being the fifth leading cause of all cancer-related deaths among women.^1^ Early stages of ovarian cancer development are extremely difficult to diagnose because of the delayed onset of symptoms, perpetuating a pattern of late-stage diagnoses in the majority of cases in the United States.^2^ Furthermore, the lack of a universal, early-detection method prevents healthcare providers from recognizing the disease until advanced metastasis has begun, resulting in most patients undergoing surgical treatment as a primary care option.^3, 4^ Depending on the extent to which the cancer has spread, patients often undergo salpingo-oophorectomy (removal of one or both ovaries and fallopian tubes, a total hysterectomy (i.e., removal of the entire uterus), a pelvic lymphadenectomy (i.e., removal of abdominal lymph nodes), and/or a omentectomy (i.e., removal of the fatty lining that encases abdominal organs). However, the current approach to oncology surgeries is oftentimes done under white-light guidance, where surgeons rely on subjective visual cues and physical palpations of the tissues of interest. This error-prone technique causes problematic negative margins, removing far excess of the healthy tissues than necessary, or may leave residual cancer cells in the body, resulting in cancer reoccurrence. ^5, 6^

Before treatment has begun, patients are commonly screened for elevated levels of cancer antigen-125 (CA-125) *via* a blood test and changes in ovary size and morphology *via* transvaginal ultrasound (TVUS).^7, 8^ While a combination of these screening tools has been used for early-detection of ovarian cancer, they suffer from poor sensitivity and false-positive results.^9^ In parallel, other imaging modalities like computed tomography (CT) and magnetic resonance imaging (MRI) have been used to detect solid tumors.^10–12^ Unfortunately, these modalities suffer from low resolution, limited sensitivity, and long scan times— all which hinder their use in a surgical setting. Fluorescence imaging has emerged at the frontier of bioimaging for its ability to address many of these issues by offering real-time feedback of the region of interest with the ability to target tissues of interest at the molecular level.^13, 14^ The utility of this technique, however, is fundamentally governed by the performance of the exogenous fluorophores, i.e. targeted fluorophores, designed to localize within malignant tissues and emit detectable photons. One such example is Pafolacianine (commercially known as CYTALUX®), which was approved by the U.S. Food and Drug Administration (FDA) in 2021 for lung and ovarian cancers.^15^ However, Pafolacianine is only useful for folate receptor alpha (FRα)-positive ovarian cancers and is limited by its fluorescence emission in the first near-infrared region (NIR-I, 700-900 nm). Indocyanine green is another popular FDA-approved fluorophore that exhibits visible light (400-700 nm) and NIR-I properties for detecting and locating ovarian sentinel lymph nodes (**Figure 1a**).^16^ Unlike the second near-infrared (NIR-II, 1000-1700 nm) region, the visible and NIR-I regions are known to exhibit increased light scattering, light absorption, and autofluorescence from surrounding biological fluids and tissues.

**Figure 1.**
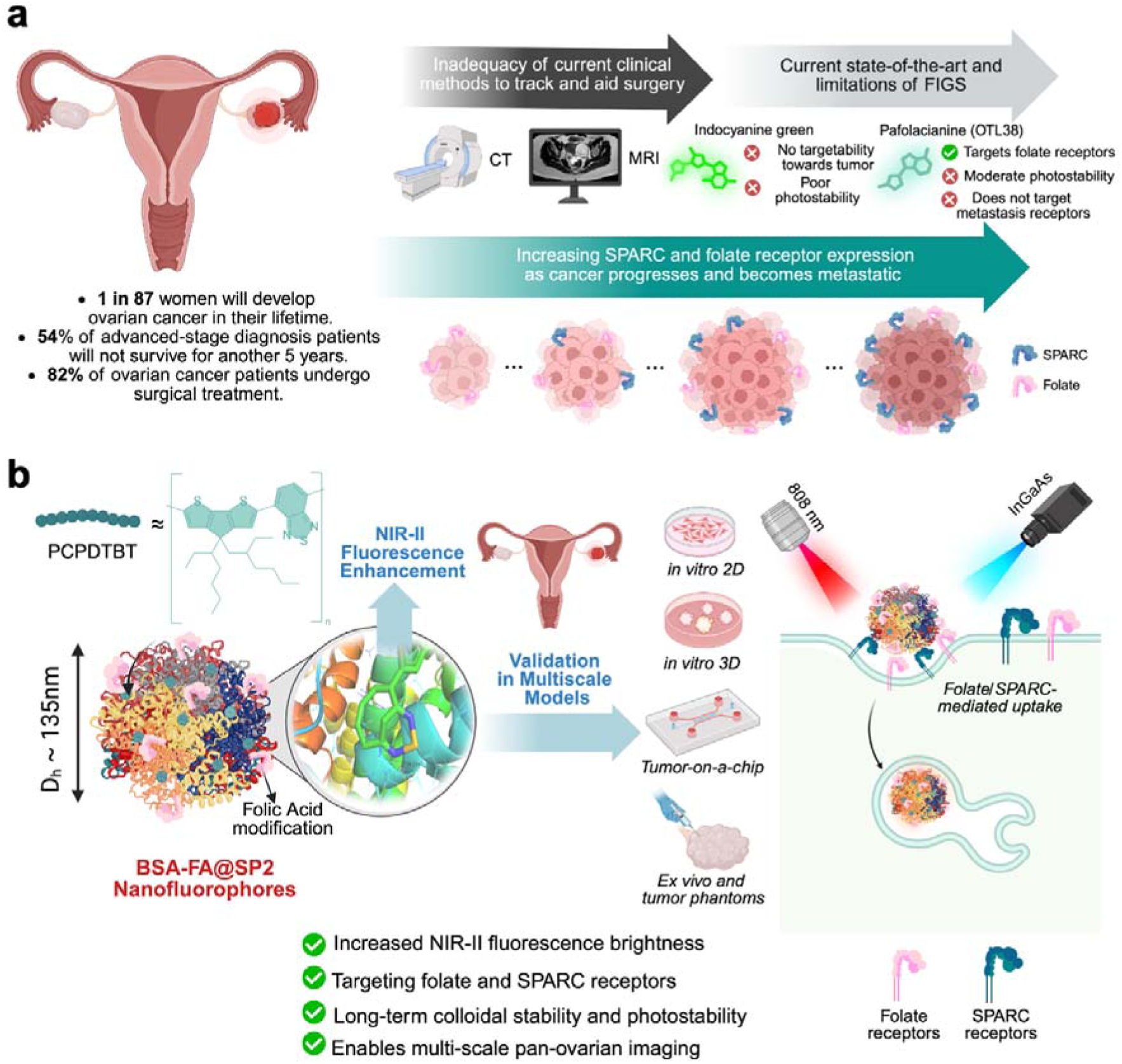
Leveraging ovarian cancer pathogenesis for the development of a universal NIR-II nanofluorophore. (a) Clinical landscape of ovarian cancer (left), including statistical outcomes and the limitations of current imaging modalities (top right).^1,2^ The schematic (bottom right) illustrates the increasing expression of SPARC and folate receptor during tumor progression and metastasis, serving as the biological rationale for targeted probe design. (b) Rationale and design of BSA-FA@SP2 nanofluorophores to amplify NIR-II fluorescence. The design utilizes a PCPDTBT polymer core encapsulated within a BSA shell, leveraging albumin-mediated amplification to significantly boost NIR-II fluorescence brightness. The probe is further modified with folic acid (FA) to enable active dual-targeting of folate (*via* FA) and SPARC (*via* BSA) receptors. The workflow details validation across multiscale models, including *in vitro* 2D, *in vitro* 3D, microfluidic tumor-on-chip models, and *ex vivo* porcine ovaries and tumor mimicking phantoms demonstrating the nanofluorophores’s efficacy in complex biological environments.

In this study, we report the rational design of a dual-receptor targeting nanofluorophores featuring synchronized NIR-I and NIR-II fluorescent signatures for high-fidelity ovarian carcinoma detection. Our platform utilizes a functional albumin matrix that leverages innate affinities for glycoprotein 60 (GP60) and secreted protein acidic and rich in cysteine (SPARC) receptors, biomarkers that are progressively upregulated during ovarian cancer metastasis.^17, 18^ To augment this intrinsic tropism, the albumin scaffold is surface-engineered via EDC-NHS mediated bioconjugation to target the overexpressed FRα receptor. By nanoconfining a representative semiconducting polymer poly[2,6-(4,4-bis-(2-ethylhexyl)-4*H*-cyclopenta[2,1-*b*;3,4-*b*′]dithiophene)-*alt*-4,7(2,1,3 benzothiadiazole)] (PCPDTBT) within the hydrophobic domains of the albumin complex, we generate a “chaperoned“ nanoflurorophore where the BSA shell serves as a robust molecular shield. This architecture effectively passivates the PCPDTBT core against non-specific protein adsorption and quenching in complex biological milieu, maintaining emission integrity in whole blood and plasma, while simultaneously exploiting the reduced photon scattering of the NIR-II window for deep-tissue visualization.

Consistent with the U.S. FDA’s recent initiatives to prioritize human-relevant, non-animal methodologies, we validated the translational utility of these nanoprobes through a systematic multiscale experimental hierarchy. Initial molecular affinity and cytocompatibility were established using *in vitro* 2D monolayers and hemocompatibility assessments, confirming negligible erythrocyte membrane disruption. Recognizing the architectural complexity of the tumor microenvironment, we progressed to *in vitro* 3D spheroids and 3D on-chip microfluidic architectures to evaluate interstitial penetration and nanoparticle transport. These microvascular models were engineered with gaps mimicking endothelial fenestrations, allowing us to rigorously quantify nanoparticle extravasation and targeting kinetics under physiological shear stress. Crucially, high cell viability was maintained across these 3D platforms, confirming the multiscale biocompatibility of the nanoprobes within complex tumor-mimicking environments. Finally, since epithelial ovarian cancer accounts for ∼90% of malignant cases^19^, we developed 3D bioprinted solid-tumor phantoms and *ex vivo* porcine ovary models to replicate the primary scattering and absorbing components of the tumor microenvironment (**Figure 1b**).

Notably, hemocompatibility assessments demonstrated negligible erythrocyte membrane disruption across physiologically relevant nanoprobe concentrations, with hemolysis remaining below 5%, comparable to negative controls. Furthermore, complement activation profiling revealed no statistically significant increase in C3a generation relative to plasma-only conditions, indicating a complement-inert, ‘stealth’ nanofluorophore behavior. These findings underscore the ability of the albumin-chaperoned architecture to minimize immune recognition and preserve systemic compatibility, key prerequisites for translational *in vivo* applications.

Building upon prior clinical successes in albumin-mediated transport and the evolving landscape of NIR-II-guided precision oncology, this chaperoned multiscale validation strategy provides an efficient, straightforward, and broadly applicable paradigm for the intraoperative visualization of complex pathologies.

## 2. Results and Discussion

### 2.1. *In Silico* Analysis of Globular Protein Interactions with PCPDTBT Polymer

Computational docking is a widely adopted *in silico* screening methodology in drug discovery, enabling rapid and accurate assessment of binding affinity and interaction geometry between small molecule ligands and protein receptors. To elucidate the binding kinetics of PCPDTBT (SP2) with potential protein chaperones, five globular proteins (bovine serum albumin, hemoglobin, lactoferrin, β-lactoglobulin, and transferrin) were evaluated as biologically relevant surfactants (**Figure 2a**). Considering the extensively delocalized π-conjugated backbone of SP2, its photophysical stability is anticipated to be influenced by non-covalent interactions within protein binding pockets.^20^

**Figure 2.**
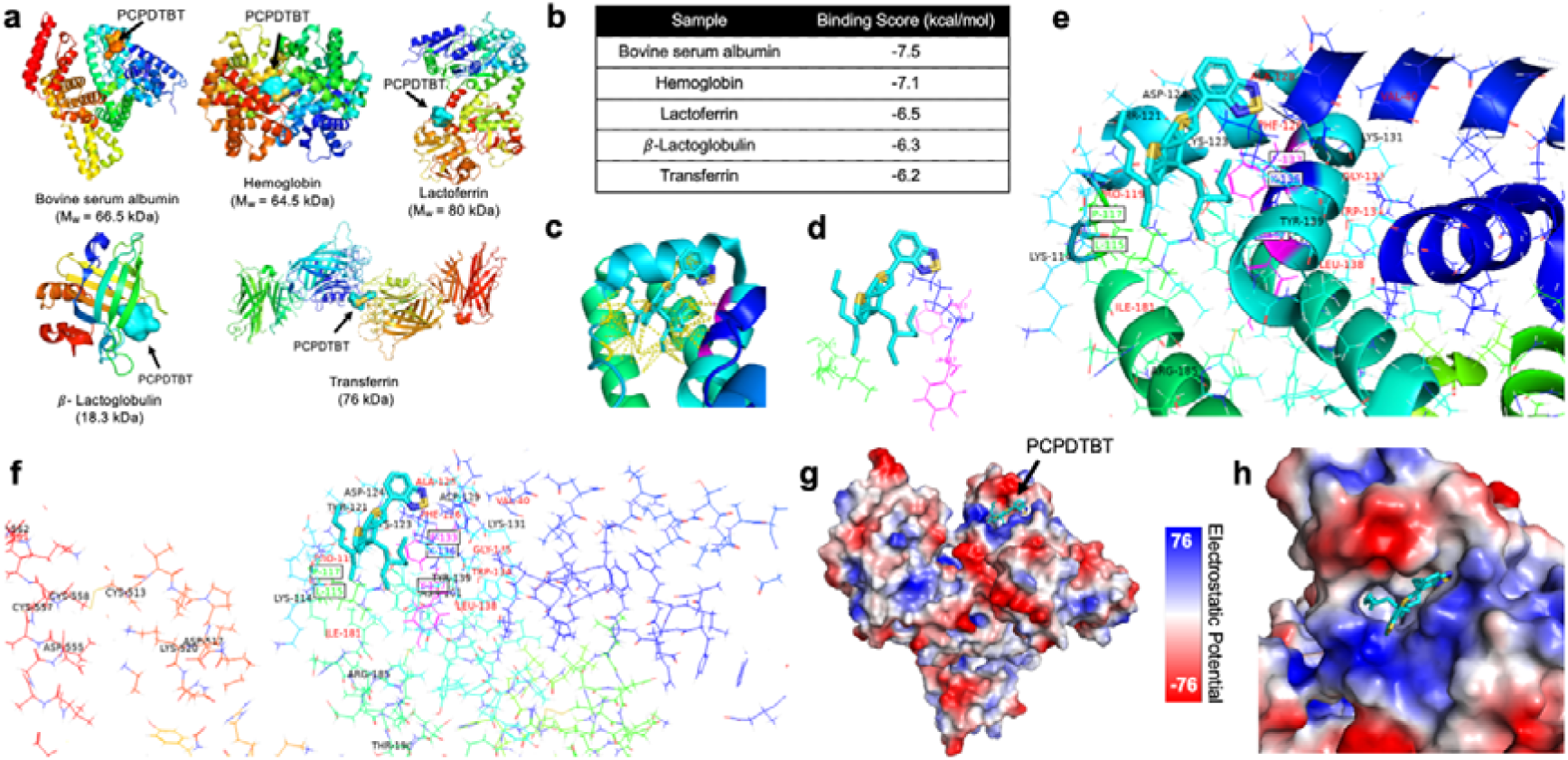
Molecular docking analyses for understanding the binding between semiconducting polymer and globular proteins. (a) 3D protein crystal structures of five globular protein candidates with a PCPDTBT monomer unit positioned at its respective binding site. (b) Corresponding average binding affinity, in kcal/mol, of PCPDTBT and each screened protein (n=3). Zoomed view to show PCPDTBT orientation within the hydrophobic cavity of BSA with (c) hydrogen bonds (yellow dashed lines) and d) π-π stacking and electrostatic interactions with surrounding amino acid residues. (e, f) Amino acids within 4.5□ of PCPDTBT. (e) Enlarged view of the binding site showing the dye and surrounding residues. (f) Representation of a selected protein segment interacting residues highlighted. Residues are categorized as hydrophilic (black), hydrophobic (red), and polar interactions (rose red box). (g, h) Electrostatic potential maps of PCPDTBT within BSA’s hydrophobic cavity. (g) Surface representation of the proteins colored by electrostatic potential. (h) Zoomed-in view of the binding pocket showing PCPDTBT within the electrostatic landscape.

Using the monomer unit of SP2 as the ligand molecule, the tertiary structure of each protein was screened to determine the binding score for SP2-protein interactions. BSA exhibited the highest binding affinity for SP2 with a docking score of -7.5 kcal/mol, corresponding to relatively strong non-covalent interactions (**Figure 2b**). Fluorescence-based titration assays between BSA and SP2 corroborated these computational predictions, yielding a dissociation constant (K_d_) of 8.067 µM (**Figure S1**).

BSA possesses well-characterized drug-binding sites known to accommodate small hydrophobic molecules, with SP2 demonstrating preferential binding to Drug Binding Site III (**Figure 2c-e**).^21–23^ Drug Binding Site III comprises a heterogenous arrangement of hydrophobic, aromatic, and charged amino acid residues that collectively contribute to both physical encapsulation and electronic stabilization of SP2.^21^ The spatial proximity of hydrophobic residues leucine and proline (L-115 and P-117) likely facilitate van der Waals and hydrophobic interactions, effectively shielding the polymer from the aqueous environment and establishing a nonpolar microenvironment (**Figure 2e, f**).^24, 25^ The aromatic amino acids phenylalanine and tyrosine (F-133 and Y-137) within the binding pocket are positioned in proximity to the π-conjugated backbone, potentially engaging in π-π stacking with the benzothiadiazole and cyclopentadithiophene moieties of SP2, thereby promoting orbital overlap and intersystem corssing (**Figure 2d-f**).^26, 27^

Electrostatic mapping was also performed using the same computational model to understand local charge distributions at the BSA-SP2 binding interface. This analysis confirmed the structural embedding of SP2 within a hydrophobic cavity that minimizes solvent exposure and thus suppresses non-radiative decay pathways and aggregation-induced quenching (**Figure 2g, h**).^28–30^ Collectively, these *in silico* analyses demonstrate the preferential binding of SP2 within a well-characterized binding site that confers both high protein affinity and enhanced photophysical stability.

### 2.2. Bioconjugation of Folic Acid to BSA Scaffolds for Enhanced Receptor Avidity

FRα is a glycosylphosphatidylinositol-anchored glycoprotein characterized by its significant overexpression in ovarian malignancies relative to its distribution in healthy tissues.^31^ The receptor features a well-characterized, open folate-binding pocket that exhibits high binding affinity for the pteridine moiety of folic acid.^32^ This distinct expression profile and the accessibility binding site render FRα a premier candidate for targeted diagnostic and therapeutic delivery systems. To facilitate active targeting of FRα-positive ovarian cancer cells, BSA was functionalized with FA via carbodiimide-mediated crosslinking chemistry (**Figure S2**). This bioconjugation strategy leverages the primary amines of the ∼20 lysine residues within the BSA’s primary sequence to form stable amide linkages with the γ-carboxyl group of FA and enable FRα targeting (**Figure S3**).^33^

The efficiency of this functionalization was corroborated *via* gel electrophoresis and mass spectrometry following extensive dialysis (**Figure S4-S6**). After the crosslinking reaction was stopped, BSA-FA complexes were purified by dialysis to remove unbound FA molecules. Removal of free FA molecules was monitored by UV-Vis absorption spectroscopy of the dialysate at 1 h and 24 h, and the final purified BSA-FA complex was also analyzed by UV-Vis spectroscopy (**Figure S4**). All albumin-based samples were subjected to sodium dodecyl sulfate polyacrylamide gel electrophoresis (SDS-PAGE), where they showed protein bands ∼66 kDa, corresponding to the molecular weight of BSA (**Figure S5**). A comparison between native BSA and BSA-FA conjugates by mass spectra revealed a molecular weight increase of ∼8797 Da. Given the molecular weight of FA (∼ 441 Da), this shift corresponds to a grafting density of approximately ∼20 FA molecules per BSA monomer (**Figure S6**). Such high-density functionalization enables the multivalent presentation of folate ligands, which is critical for enhancing receptor avidity while maintaining the protein’s native structural integrity and solubility. Upon binding the FRα, the resulting BSA-FA conjugate triggers membrane invagination, facilitating cellular entry *via* receptor-mediated endocytosis.^34,35^ Robust amide connectivity ensures the integrity of the complex during systemic circulation, preventing premature ligand dissociation and ensuring high-fidelity delivery to the intracellular compartment.

### 2.3. Synthesis, Physicochemical and Optical Characterization of BSA-FA@SP2 Nanofluorophores

To systematically evaluate colloidal stability, photophysical properties, and targeted ovarian cancer imaging efficacy, three distinct NP formulations were developed and characterized in this study. A benchmark control was synthesized using 1,2-distearoyl-sn-glycero-3-phosphoethanolamine-poly(ethylene glycol) (DSPE-PEG2k), a clinically approved amphiphilic polymer widely employed in FDA-approved nanomedicines such as Doxil^®^.^36^ In parallel, BSA and BSA-FA NPs were utilized as biomolecular scaffolds to encapsulate the SP2 fluorophore within the protein’s surface-accessible hydrophobic pockets domains. These three NP platforms were designed for the rigorous assessment of: (i) the ability to form stable self-assembly and optically active nanofluorophores suitable for bioimaging applications, and (ii) the quantitative impact of FA-conjugation on FRα-mediated endocytosis (**Figure 3a**).

**Figure 3.**
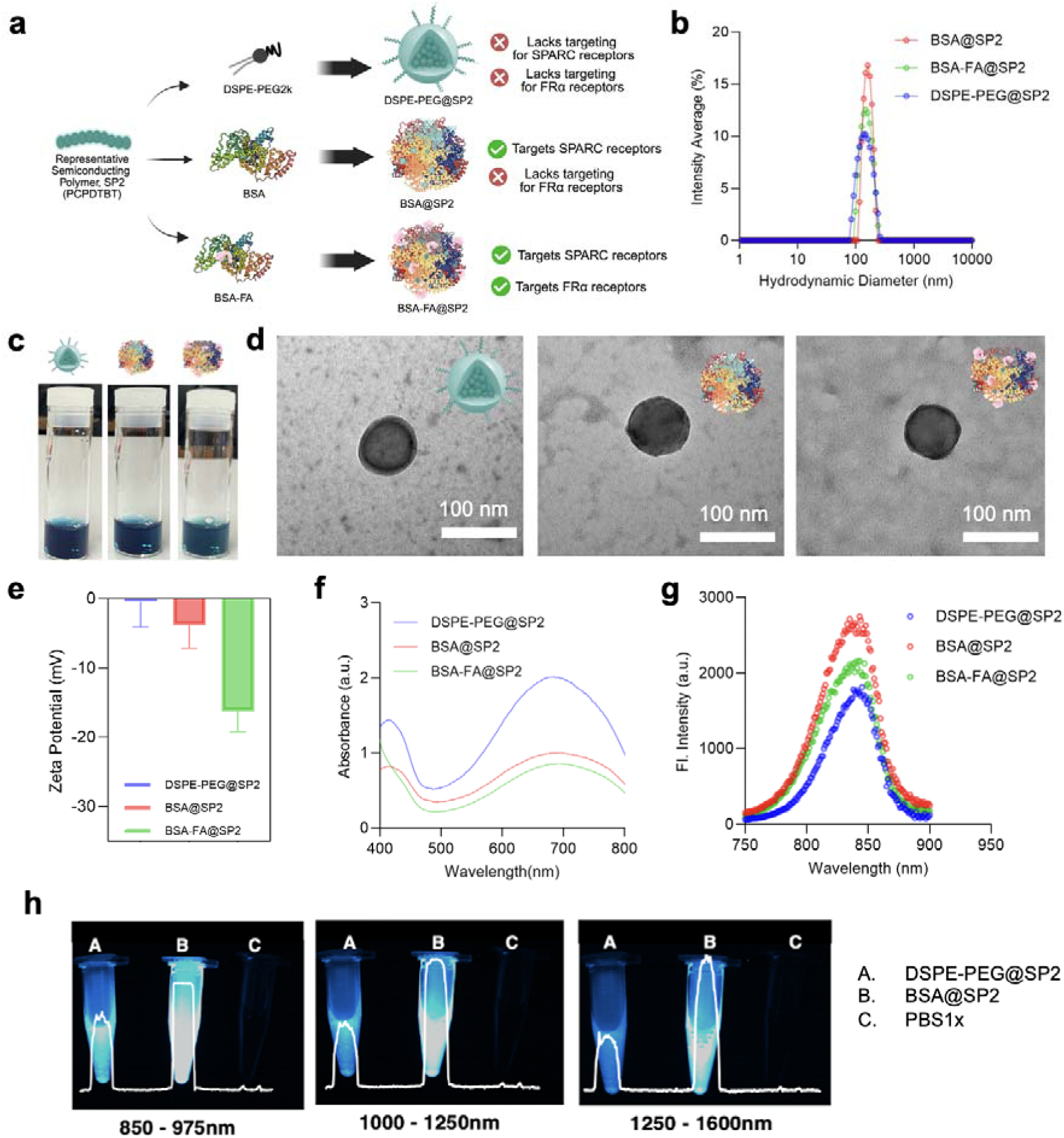
Preliminary physicochemical and optical characterization of experimental NPs. (a) Schematic illustration of NP designs (DSPE-PEG@, BSA@, and BSA-FA@SP2) for active targeting of ovarian cancer-associated biomarkers. (b) Dynamic light scattering plot of hydrodynamic diameter intensity average of DSPE-PEG@, BSA@, and BSA-FA@SP2. (c) White light images of DSPE-PEG@, BSA@, and BSA-FA@SP2 samples at 100 µg/ mL. (d) Representative TEM images of DSPE-PEG@, BSA@, and BSA-FA@SP2 (left to right). (e) Zeta-potential, (f) UV-VIS-NIR absorbance spectra, and (g) fluorescence spectra (λ_ex_: 645 nm) of DSPE-PEG@, BSA@, and BSA-FA@SP2. (h) NIR-II images of (A) DSPE-PEG@SP2, (B) BSA@SP2, and (C) PBS1x within three emission windows following excitation at 808 nm.

All BSA-based nanofluorophores were formulated using 100 µM of BSA to employ ideal colloidal and optical properties (**Figure S7**). The hydrodynamic diameter of the formulations was interrogated *via* dynamic light scattering, and BSA-FA@SP2 nanofluorophores exhibited a hydrodynamic diameter (D**_h_**) of 180 nm, representing a marginal increase of approximately 30 nm compared to unmodified BSA@SP2 nanofluorophores (**Figure 3b**). The low polydispersity index (PDI) of 0.10 confirms a monodisperse BSA-FA@SP2 population, and the colloidal stability in physiologically-relevant environments (37°C, plasma) are essential for *in vivo* transport (**Figure S8, S9**). All three nanofluorophore designs (DSPE-PEG, BSA@, and BSA-FA@SP2) (**Figure 3c**) resulted in NP populations with uniform, spherical morphologies (**Figure 3d**).

Surface chemistry modifications were further validated by zeta potential measurements; BSA-FA@SP2 nanofluorophores displayed a more negative surface charge (-16.2 ± 2.96 mV) compared to BSA@SP2 (-3.77 ± 3.39 mV), a shift consistent with the introduction of ionizable α-carboxyl groups from the conjugated FA moieties (**Figure 3e**). Bicinchoninic acid (BCA) assays confirmed that protein (i.e. BSA) was exclusively present only in the BSA-based formulations (BSA@SP2 and BSA-FA@SP2). In contrast, the DSPE-PEG@SP2 control showed no detectable protein content, verifying the compositional fidelity of each formulation (**Figure S10**).

The photophysical properties of the encapsulated SP2 were evaluated using UV-Vis absorption and fluorescence spectroscopy. All three formulations maintained consistent absorbance maxima and fluorescence emission profiles (**Figure 3f, g**), suggesting that the electronic ground state of the SP2 guest remains unperturbed by the different encapsulation matrices. Notably, both BSA-based nanofluorophores (BSA@SP2 and BSA-FA@SP2) exhibited marked enhanced in radiant emission intensity relative to DSPE-PEG@SP2 control (**Figure 3h**). This protein-induced fluorescence enhancement is attributed to the entrapment of the SP2 polymer within the BSA hydrophobic pockets, which restricts intramolecular rotation and effectively suppresses non-radiative decay pathways.^37, 38^ Furthermore, the protein scaffold serves as a steric shield, protecting the fluorophore from solvent-induced quenching, ensuring NIR-II signal enhancement. ^37, 38^ This observation is consistent with previously reported BSA-fluorophore interactions, where the protein microenvironment modulates the photophysical properties of encapsulated payloads to favor radiative transitions over non-radiative dissipation.^38^

### 2.4. Nanofluorophores Biocompatibility and Cellular Uptake in *In Vitro* 2D Models

To evaluate the cellular internalization of BSA-FA@SP2 nanofluorophores in ovarian cancer cells within *in vitro* 2D models with documented FOLR1 and SPARC expression, we assessed the platform across four ovarian cancer cell lines, including three human-derived models (OVCAR3, OVCAR8, SKOV3) and one murine line (ID8).^39–41^ Mechanistic and time-resolved uptake studies were primarily conducted in SKOV3 and OVCAR8 due to their documented transcriptomic co-expression of SPARC and FRα (**Figure 5a**). Complementary validation of biocompatibility and cellular uptake in OVCAR8, ID8, OVCAR3 is provided in the Supporting Information (**Figure S11-S12**). Initial cytotoxicity screening demonstrated that BSA-FA@SP2 exhibits negligible cytotoxicity at concentrations up to 200 µg/ mL in SKOV3 (**Figure 5b**), establishing a safe and biocompatible working range for subsequent 2D uptake studies. Comparative uptake analysis revealed that BSA-FA@SP2 nanofluorophores achieved significantly higher cellular internalization than both DSPE-PEG@SP2 and BSA@SP2 nanofluorophores across both SKOV3 and OVCAR8 cell lines (**Figure 5c, d, S13**). This enhanced accumulation is consistent with increased targeting efficacy arising from dual receptor engagement, leveraging both SPARC-associated albumin binding and folate receptor-mediated endocytosis.

**Figure 5.**
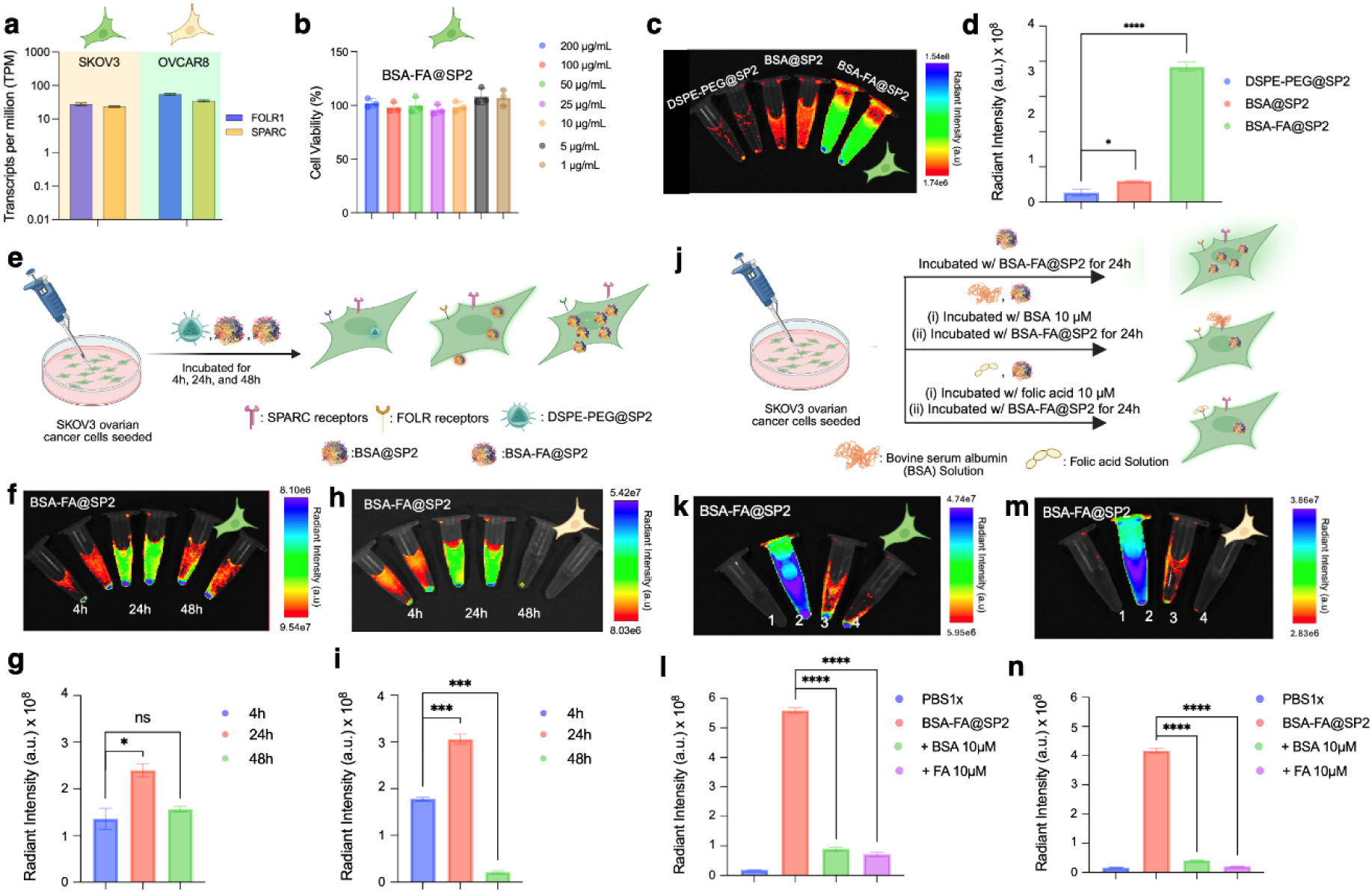
Evaluation of BSA-FA@SP2 nanofluorophores using *In Vitro* 2D Models. (a) RNA sequencing data confirming the biomarker expression levels of FOLR1 and SPARC genes in SKOV3 and OVCAR8 tumor cells. (b) SKOV3 cell viability following BSA-FA@SP2 treatment [1-200µg/mL]. (c) IVIS fluorescence images of SKOV3 cultures after 24h treatment with DSPE-PEG@, BSA@, and BSA-FA@SP2 nanofluorophores, with (d) corresponding radiant intensities. (e) Schematic diagram showing experimental designs for time-dependent uptake studies. (f,h) IVIS fluorescence images of cellular uptake of BSA-FA@SP2 in (f) SKOV3 and (h) OVCAR8, with (g,i) corresponding radiant intensities. (j) Schematic diagram showing experimental designs for inhibition studies. (k, m) IVIS fluorescence images of cell uptake of BSA-FA@SP2 in (k) SKOV3 and (m) OVCAR8 following treatment with media (control: microcentrifuge tube 2), SPARC-blocking solution (10µM, microcentrifuge tube 3), and FOLR-blocking solution (10µM, microcentrifuge tube 4). microcentrifuge tube 1 serves as a cell media control, with (l,n) corresponding radiant intensities. All IVIS images were collected with λ_ex_: 675nm and λ_em_: 840nm.

To determine time-dependent nanofluorophore uptake patterns, both cell lines were treated with 100 µg/mL BSA-FA@SP2 for 4, 24, and 48 hours (**Figure 5e**). IVIS images of harvested cell pellets demonstrated that BSA-FA@SP2 exhibited maximal cellular internalization at 24 hours (**Figure 5f, h**), with fluorescent intensity significantly diminished at 48 hours (**Figure g, i**). The peak uptake observed at 24 hours likely reflects a steady-state equilibrium between active receptor-mediated endocytosis and intracellular trafficking processes.^42^ At this time point, sufficient internalization has occurred while minimizing endocytic pathway saturation and subsequent nanofluorophore exocytosis.^43^

To elucidate the contribution of surface receptors to the active uptake mechanism of BSA-FA@SP2, competitive inhibition studies were performed by pre-treating cells with 10 µM SPARC- and folate receptor-blocking solutions (**Figure 5j**). IVIS images revealed that receptor blockade significantly attenuated BSA-FA@SP2 uptake (**Figure 5k, m**), with uninhibited cells displaying approximately 4-fold higher fluorescent intensity compared to pre-treated counterparts (**Figure 5l, n**). These findings confirm that BSA-FA@SP2 internalization occurs primarily through SPARC- and FRα- dependent pathways, validating the dual-targeting design strategy.

### 2.5. Nanofluorophore Biocompatibility and Cellular Uptake in 3D Tumor Spheroids

While 2D monolayer cultures provide valuable insight into receptor-specific binding and initial cytotoxicity, they fail to recapitulate the structural and physiological complexity of solid tumors. In particular, the dense cellular architecture, extracellular matrix deposition, and diffusion-limited transport within tumors present significant barriers to nanoparticle penetration and uniform distribution. Transitioning to 3D multicellular tumor spheroids enables a more physiologically relevant evaluation of nanofluorophore behavior, including interstitial diffusion, depth-dependent uptake, and retention within tumor-like microenvironments.^44, 45^ This model is critical to determine whether BSA-FA@SP2 nanofluorophores can effectively penetrate beyond the proliferative outer layers and maintain targeting specificity throughout the spheroid core. Furthermore, assessing cell viability within this architecture provides insight into the biocompatibility of the nanofluorophores under conditions that more closely mimic in vivo tumor organization and metabolic gradients.

While SKOV3 and OVCAR8 were prioritized in Section 2.4 for mechanistic uptake studies due to their high co-expression of SPARC and FRα as observed from RNA sequencing (**Figure 5a**), 3D spheroid evaluation was conducted using OVCAR3 and ID8 models to assess nanofluorophore performance in more heterogeneous tumor architectures and to incorporate a murine system for translational relevance. Ovarian cancer spheroids were initiated with 1000 OVCAR3 or ID8 cells and allowed to mature for 4 days before a 24-hour NP exposure (**Figure 6a**). Representative phase contrast micrographs demonstrate minimal cell death after 24 hours of NP exposure for OVCAR3 (**Figure 6b**) and ID8 (**Figure 6d**) spheroids. Viability was visualized microscopically by the maintenance of spheroid structural integrity in untreated controls as well as NP-treated spheroids. Orthogonal quantification of viability using a metabolic readout also indicated minimal changes with NP treatment across both spheroid models. OVCAR3 spheroids maintained high viability following 24 hours of nanoparticle exposure, with 25 µg/mL BSA-FA@SP2 inducing only a slight decrease in cell viability (14.13%, ** <0.0001, two-way ANOVA, **Figure 6c**). ID8 spheroids had no significant decreases in viability across all 24-hour nanoparticle treatments (two-way ANOVA, **Figure 6e**). Negligible cytotoxicity was similarly observed after 48 hours of exposure in OVCAR3 and ID8 spheroids (**Figure S14**). Compared to the clinical standard DSPE-PEG@SP2, BSA@SP2 and BSA-FA@SP2 demonstrate increased OVCAR3 and ID8 spheroid uptake after 48 hours of exposure to 25 µg/mL nanofluorophores, indicated by increased radiant intensity of spheroid lysates (**Figure 6f, g**). Spheroid uptake of nanofluorophores across varying concentrations over 24 and 48 hours of exposure show similar uptake trends (**Figure S15**).

**Figure 6.**
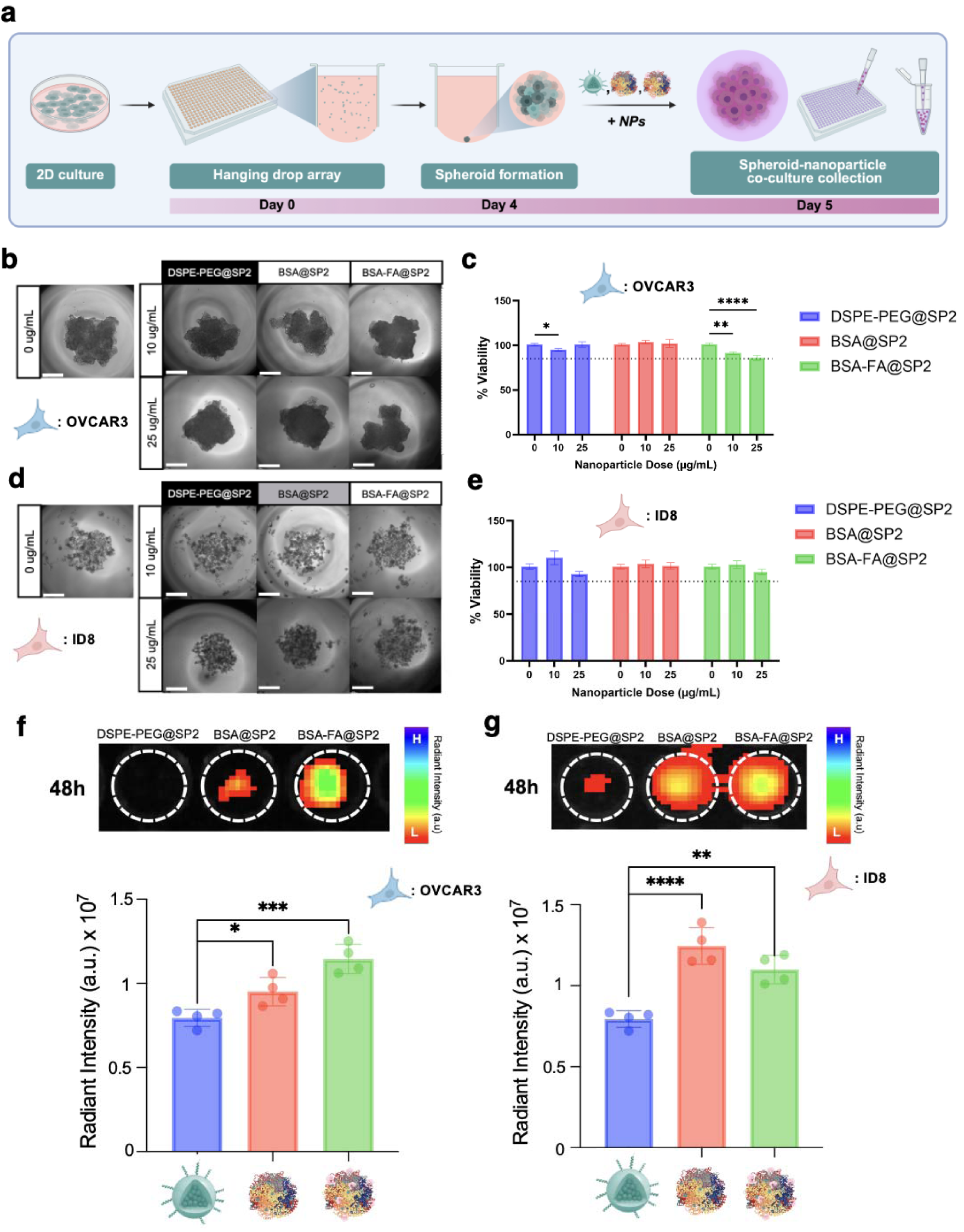
Evaluation of BSA-FA@SP2 using 3D *In Vitro* Spheroid Models. (a) Schematic illustration of 3D spheroid formation and treatment protocol. Phase contrast micrographs of control or nanoparticle treated spheroids, and quantification of their corresponding viability following 24hr treatments in OVCAR3 (b, c) or ID8 (d, e) spheroids. IVIS images (top) and corresponding fluorescent intensity (bottom) of (f) OVCAR3 and (g) ID8 spheroid lysates following 48-hour NP treatments.

### 2.6. Flow-Dependent Nanofluorophore Transport and Uptake in Tumor-on-a-Chip Microfluidic Models

While 2D cultures and static 3D spheroids provide initial insights into cellular affinity, they fail to account for the hemodynamic forces and mass transport limitations inherent in the complex tumor microenvironment.^46^ Moving to a flow-based tumor-on-a-chip model is essential to evaluate whether the BSA-FA@SP2 nanofluorophore can maintain its targeting integrity under physiological shear stress and successfully extravasate through “leaky” endothelial barriers to reach the tumor interstitium.^47, 48^ This platform bridges the gap between *in vitro* assays and *in vivo* performance by providing a more rigorous assessment of the nanoparticle’s transport kinetics and receptor-mediated uptake within a biomimetic architecture.

To evaluate these dynamics, our 3D-bioprinted perfusable platform featured a microfluidic channel separated from a 3D SKOV3 tumor cluster by a semi-permeable interface mimicking endothelial fenestration (**Figure 7a**).^49^ Based on our preliminary *in vitro* 2D screening across a panel of ovarian cancer cell lines, including SKOV3, OVCAR8, OVCAR3, and murine ID8 cells, SKOV3 was selected for on-chip validation as dual-receptor-positive model. Specifically, its robust expression of both folate receptors and SPARC effectively captures the synergistic targeting potential of our BSA-FA@SP2 nanofluorophores systems in a highly invasive microenvironment.

**Figure 7.**
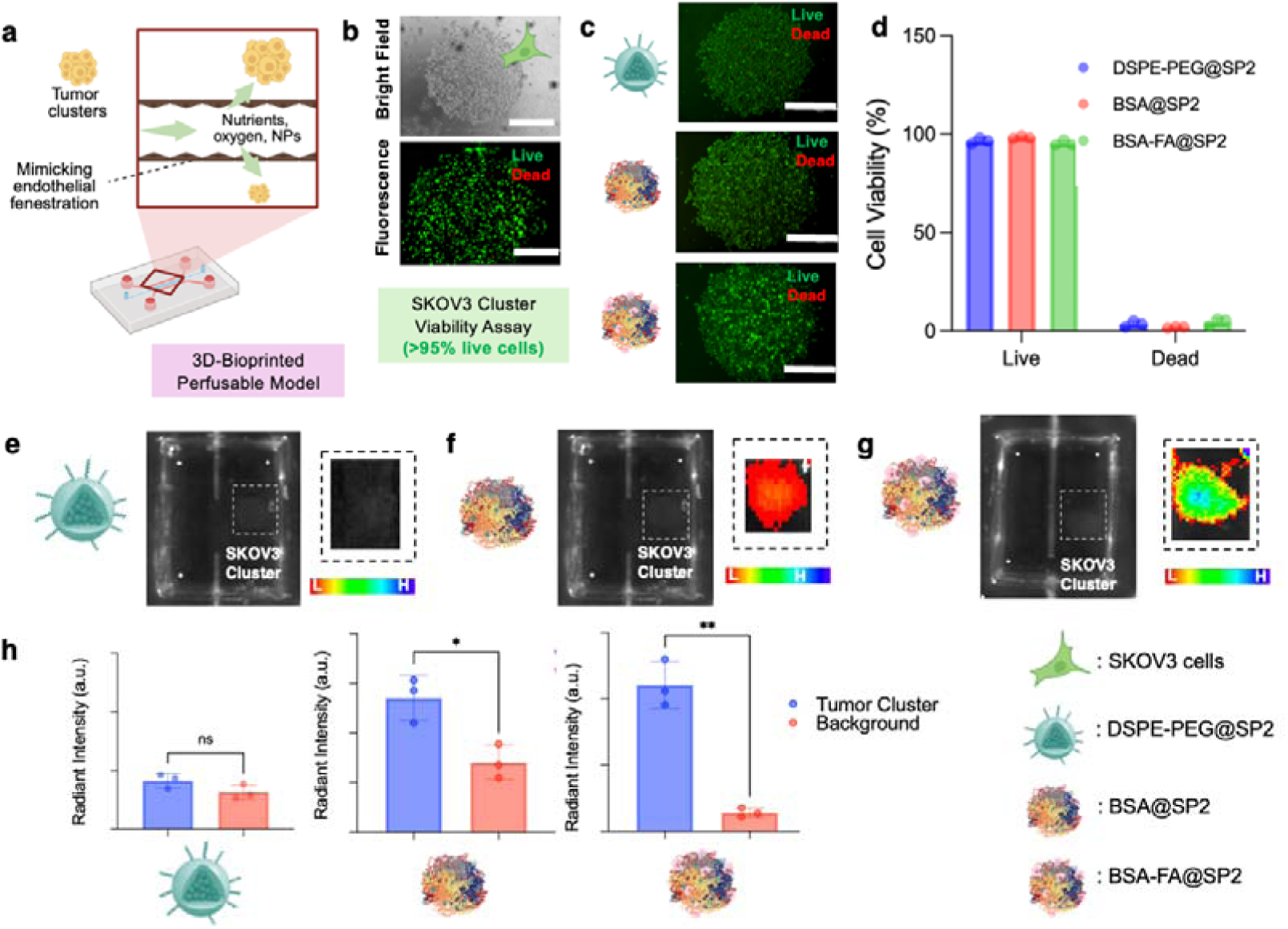
Assessment of BSA-FA@SP2 Nanofluorophore Transport and Cellular Uptake in a Tumor-on-a-Chip Microfluidic Model Mimicking Endothelial Leakiness. (a) Schematic of the on-chip platform featuring a microfluidic channel mimicking leaky vasculature and a tumor cluster. (b) Representative bright-field (top) and live/dead fluorescence (bottom) images of SKOV3 cell clusters. (c) Live/dead assay fluorescence images of SKOV3 clusters following 36h perfusion of nanofluorophore treatments with (d) corresponding cell viability quantification using a MATLAB script (See Experimental Methods), n = 6. (e-g) IVIS images of SKOV3 clusters after 36h perfusion with (e) DSPE-PEG@SP2, (f) BSA@SP2, and (g) BSA-FA@SP2 with (h) respective radiant intensities, n = 6. All IVIS images were collected with λ_ex_: 675nm and λ_em_: 840nm.

Bright-field and fluorescence microscopy confirmed the successful formation of dense, spherical SKOV3 clusters within the GelMA bioink (**Figure 7b**). Initial live/dead assays demonstrated high viability (∼99%), confirming that the 3D-bioprinting process and the covalently crosslinked GelMA hydrogel matrix provide a biocompatible environment conducive to ovarian cancer cell growth. The cytotoxicity of the nanofluorophore formulations was assessed following 36 hours of continuous perfusion. Live/dead fluorescence imaging (**Figure 7c**) and subsequent quantification *via* a custom MATLAB script (**Figure 7d**) revealed that all formulations, including DSPE-PEG@SP2, BSA@SP2, and BSA-FA@SP2, maintained high cell viability. The lack of significant “dead” cell populations across all treatment groups confirms that these SP2-based organic nanofluorophores are highly biocompatible and suitable for prolonged diagnostic imaging in 3D biological environments.

The targeting efficacy of the functionalized nanofluorophore was further evaluated through IVIS analysis of the chip. While DSPE-PEG@SP2 nanofluorophores showed negligible accumulation (**Figure 7e, h**), the BSA-FA@SP2 nanofluorophores exhibited significantly higher radiant intensity and distinct localization within the SKOV3 clusters (**Figure 7g, h**). This enhanced accumulation is attributed to a synergistic dual-targeting mechanism: the BSA coating facilitates initial transport and interstitial localization through its affinity for SPARC overexpressed in the tumor microenvironment, while the folic acid ligands promote high-affinity, receptor-mediated endocytosis by the folate receptors overexpressed in SKOV3 cells.^39, 50^ The intermediate performances of BSA@SP2 nanofluorophores (**Figure 7f, h**) compared to PEGylated nanofluorophores further supports the role of albumin-SPARC interactions in improving tumor-cluster retention. These results demonstrate that the BSA-FA@SP2 nanofluorophore effectively navigates simulated vascular barriers and achieves specific, high-contrast targeting in a complex, biomimetic 3D environment, validating its potential for precise NIR-II ovarian cancer imaging.

### 2.7. Evaluating Optical Performances of BSA-FA@SP2 Nanofluorophores in Simulated Biological Environments

To evaluate whether biologically relevant media affect the optical performance of BSA-FA@SP2 nanofluorophores, we measured its NIR-II fluorescence following incubation with fetal bovine serum (FBS), plasma, and whole blood. While these complex biological fluids contain growth factors, lipids and diverse proteins that typically induce intermolecular quenching or non-specific aggregation, BSA-FA@SP2 nanofluorophores maintained NIR-II fluorescence intensities comparable to those in buffer alone (Figure 8a).^51–53^ Such observations indicate minimal quenching under simulated *in vivo* like environments. The high degree of emission integrity suggests that the BSA shell acts as a robust molecular shield^54, 55^, protecting the PCPDTBT core from quenching interactions while maintaining the albumin-mediated amplification behavior of NIR-II signal.^37, 38, 55^

**Figure 8.**
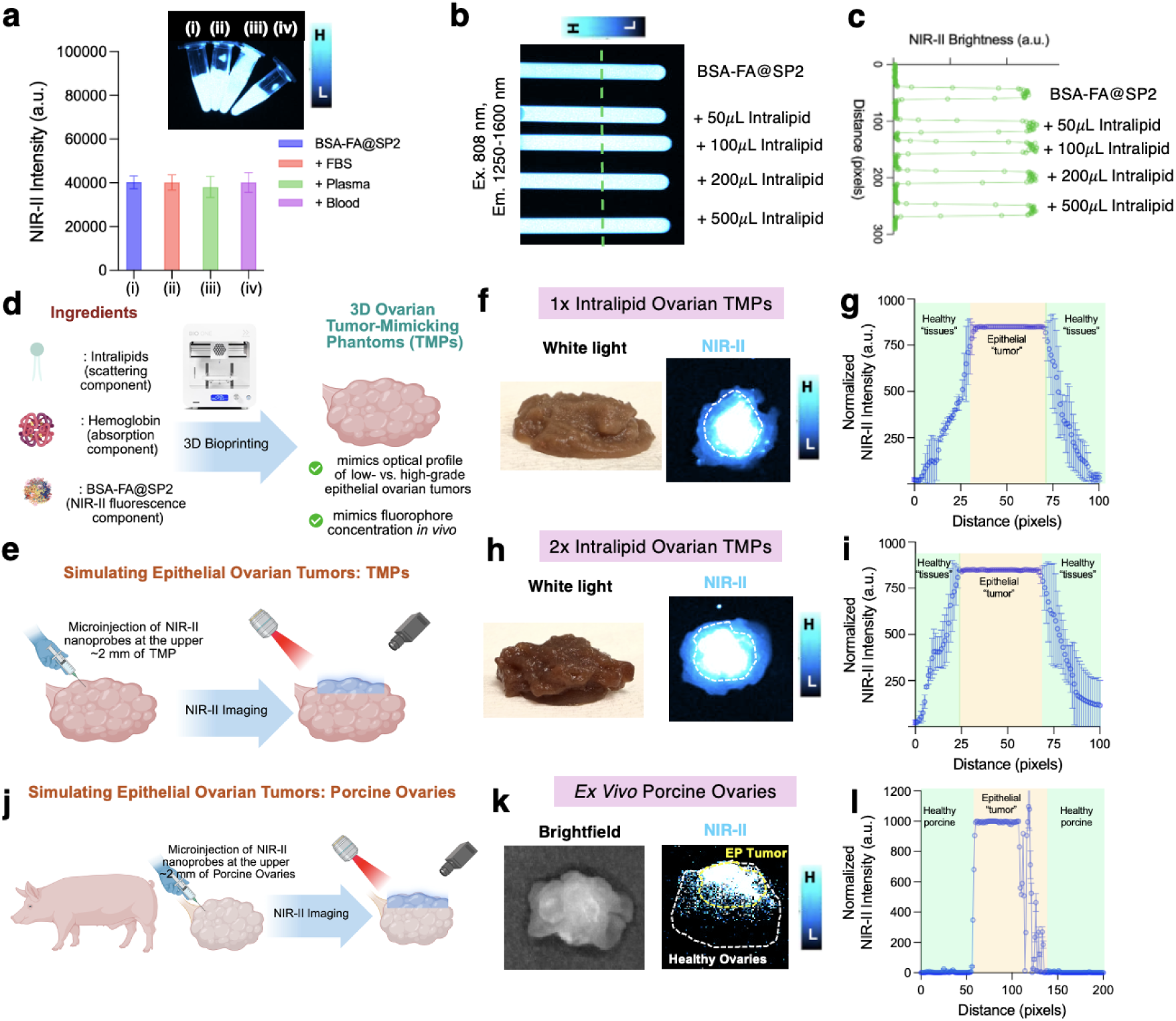
Optical Performance of BSA-FA@SP2 Nanofluorophores for Intraoperative Guidance. (a) NIR-II fluorescence intensity of BSA-FA@SP2 in (i) PBS1x solution, (ii) fetal bovine serum (FBS), (iii) plasma, and (iv) whole blood. (b) NIR-II fluorescence image of BSA-FA@SP2 with increasing intralipid content, with (c) corresponding NIR-II brightness. (d, e) Schematic illustration of the formulation and design of 3D bioprinted tumor-mimicking phantoms, including nanoprobe composition (d) and microinjection strategy (e). (f, h) Representative white light (left) and NIR-II fluorescent images (right) of the phantoms, with corresponding (g, i) NIR-II fluorescent intensity of healthy regions and epithelial tumor regions at varying lipid content. (j) *Ex vivo* porcine ovary model following microinjection of BSA-FA@SP2 to simulate epithelial tumors, with (k) white light (left) and NIR-II fluorescent images and (l) corresponding NIR-II fluorescent intensity. All IR VIVO images were acquired at λ_ex_: 808 nm and λ_em_: 1250 – 1600 nm

Furthermore, because lipid-rich tissues often degrade optical signals through significant light scattering^56, 57^, a phenomenon pronounced in the visible and NIR-I ranges, we assessed nanofluorophores’ performance in the presence of Intralipid as a tissue-mimicking scattering agent. Consistent with the reduced scattering properties in the NIR-II window^58, 59^, BSA-FA@SP2 nanofluorophores showed negligible loss of radiant intensity even at high intralipid concentrations (**Figure 8b, c**). These results collectively demonstrate that BSA-FA@SP2 nanofluorophores retains robust NIR-II emission in complex biological fluids and remains insensitive to lipid-induced scattering, underscores its suitability for high-fidelity, deep-tissue imaging in physiologically relevant environments.

### 2.8. NIR-II Fluorescence Imaging of Microinjected 3D Bioprinted Phantoms and *Ex Vivo* Porcine Ovaries Modeling Epithelial Tumors

Following the U.S. Food and Drug Administration’s announcement to phase out conventional animal testing in favor of more human-relevant models^60^, increasing attention has been directed toward tumor phantoms that more accurately replicate the optical environment of human tissues.^61, 62^ Because epithelial ovarian cancers accounts for approximately 90% of all malignant ovarian cases and frequently manifests as superficial metastases, developing high-fidelity models of these lesions is critical for validating intraoperative probes. Here, we employed a solid-tumor phantom formulation containing Intralipid and hemoglobin to mimic the primary scattering and absorbing components of the tumor microenvironment (**Figure 8d**). To simulate the localized nature of epithelial lesions, we used a sequential validation strategy: an ovarian tissue model was 3D-printed using the optimized phantom recipe to establish a physiologically relevant optical background (**Figure 8d**). This was followed by a microinjection of BSA-FA@SP2 nanofluorophore to create a defined fluorescent inclusion (**Figure 8e**).

By systematically increasing the Intralipid concentration from 1x to 2x, we were able to tune the optical and physical properties of the matrix to simulate progressive epithelial tumor stages. This increase in scattering was accompanied by a visibly more rigid phantom morphology, effectively mimicking the increased density and structural stiffness characteristic of advanced tumors. Across both cases, NIR-II imaging provided clear contrast between the surrounding “healthy” phantom matrix and the localized BSA-FA@SP2 nanofluorophore accumulation, enabling straightforward delineation of the simulated tumor region (**Figure 8f-i, S16**). Tissue-stacking studies demonstrated the ability of these BSA-based nanofluorophores to penetrate through porcine muscle, fat, and skin tissues within a similar simulate tumor environment (**Figure S17 – S19**).

To extend these findings to biological tissue, we performed an *ex vivo* validation using porcine ovaries. BSA-FA@SP2 nanofluorophores was microinjected ∼2 mm below the ovarian surface to mimic a superficial epithelial lesion (**Figure 8j**). Consistent with the phantom results, NIR-II imaging showed distinct localized signal at the injection site with minimal background interferences from the surrounding healthy tissues (**Figure 8k, l**). While *ex vivo* model have inherent limitations in recapitulating dynamic *in vivo* perfusion, they provide a critical testbed for assessing signal attenuation and contrast in real tissue architecture. Collectively, these studies demonstrate that BSA-FA@SP2 nanofluorophores enables robust NIR-II contrast and reliable tumor delineation, supporting its potential utility for intraoperative visualization of superficial epithelial ovarian lesions.

### 2.9. Hemocompatibility Assessments of BSA-FA@SP2 Nanofluorophores

A key requirement for blood-contacting biomaterials is preserving erythrocyte (red blood cell, RBC) membrane integrity. RBC rupture (hemolysis) can lead to reduced oxygen delivery and elevated level of free hemoglobin in circulation, which may cause systematic toxicity.^63, 64^ Here, BSA-FA@SP2 was incubated with paced RBCs at physiological temperature to quantify hemolysis (**Figure 9a**). After one hour, all BSA-FA@SP2 concentrations tested (1-100 µg/mL) caused minimal membrane disruption (**Figure 9b**), comparable to the PBS1x negative control (**Figure 9c, d**). Consistent with these observations, hemolysis remained below 5% for all BSA-FA@SP2 treatments, whereas DI water produced near-complete hemolysis (∼100%) (**Figure 9e**).

**Figure 9.**
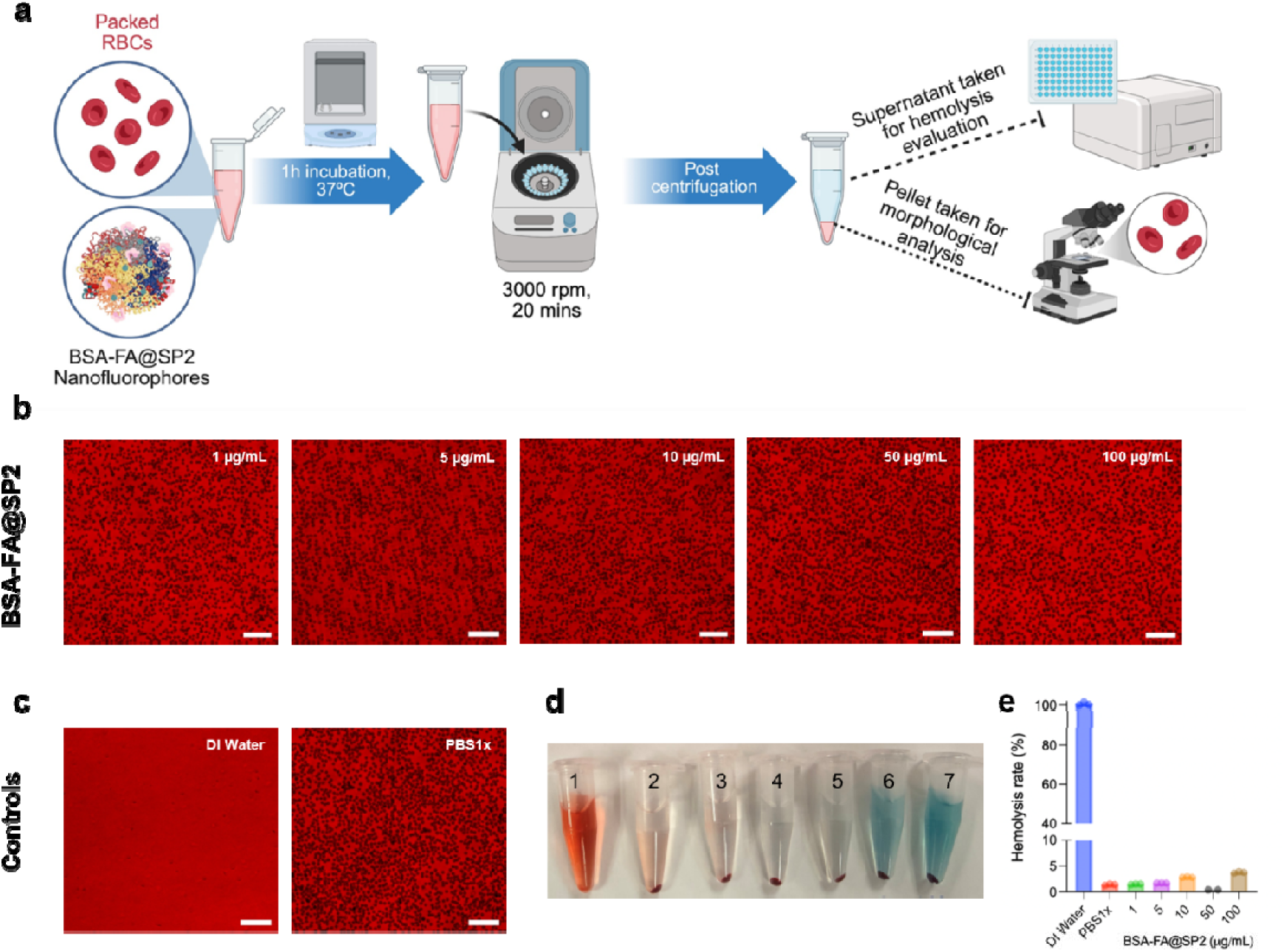
Hemocompatibility Assessment of BSA-FA@SP2. (a) Schematic illustration of the hemolysis assay workflow used to evaluate RBC compatibility following treatment with BSA-FA@SP2. Representative microscope images (200x magnification) of RBC pellets after incubation with (b) BSA-FA@SP2 and (c) control groups. Scale bar = 100 µm. (d) Image of RBC pellet formation in microcentrifuge tubes after treatment with (1) DI water (positive control), (2) PBS1x (negative control), and BSA-FA@SP2 at (3) 1, (4) 5, (5) 10, (6) 50, and (7) 100 µg/mL. (e) Percent hemolysis for all groups, calculated from the absorbance of released hemoglobin at 540 nm.

### 2.10. Complement Activation Profiling Reveals Stealth Behavior of BSA-FA@SP2 Nanofluorophores

To evaluate the immunological compatibility of BSA-FA@SP2 nanofluorophores in a physiologically relevant biological milieu, we quantified complement activation using a C3a-specific ELISA assay following incubation with plasma (**Figure 10a**). C3a is a well-established anaphylatoxin generated upon proteolytic cleavage of complement component C3 and serves as a sensitive biomarker for early stage complement activation. Upon activation, C3 cleavage yields C3b, which mediates opsonization, and C3a, which diffuses into the surrounding fluid phase and can trigger inflammatory signaling cascades through mast cell and basophil activation (**Figure 10b**).^65^ The C3a ELISA used here detects the captured human C3a through a sandwich immunoassay with a colorimetric readout at 450 nm proportional to C3a concentration. A linear calibration curve (R^2^ = 0.93) confirmed the assay’s quantitative reliability across the tested concentration range (**Figure 10c**).

**Figure 10.**
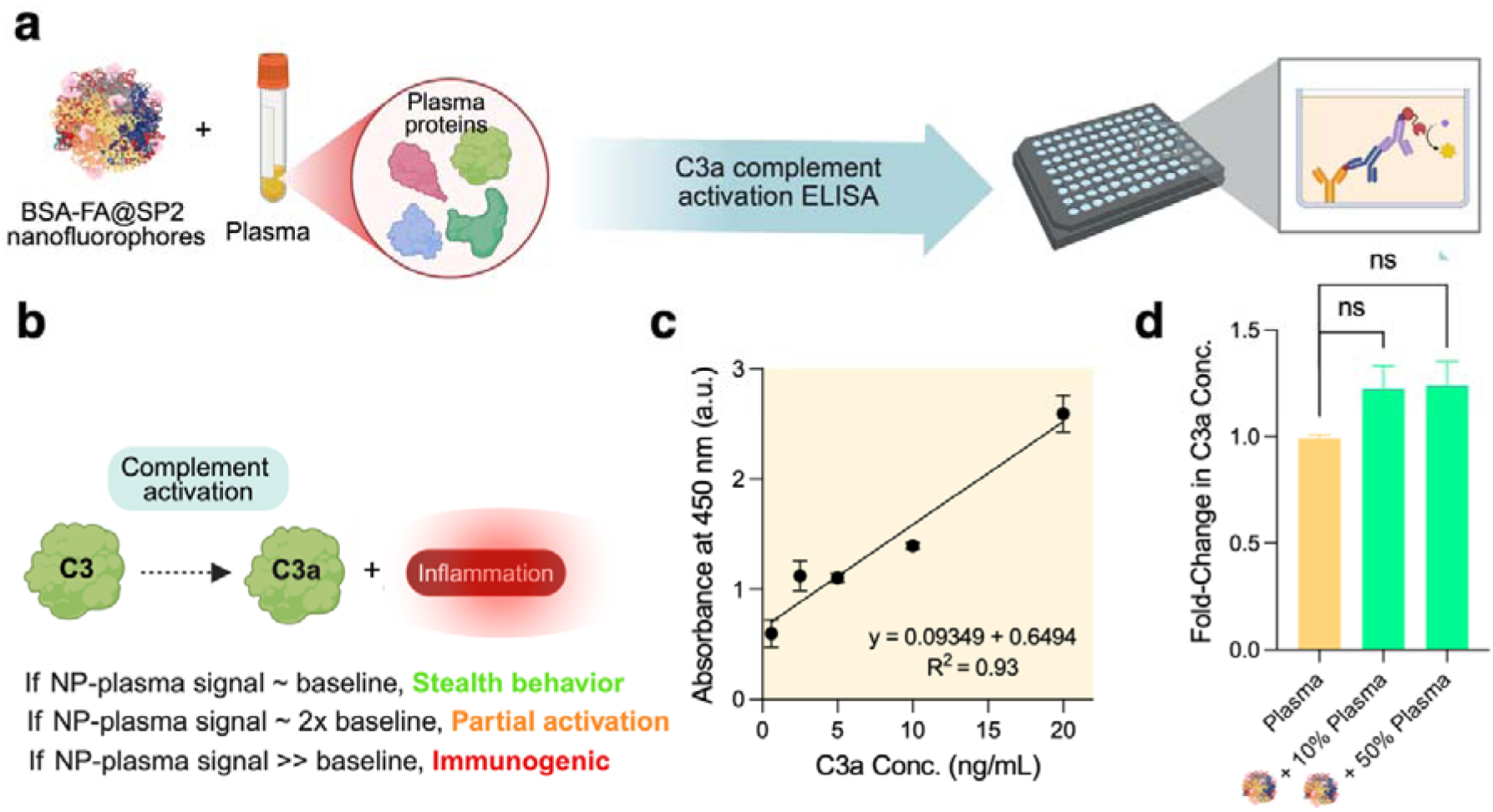
Evaluation of complement activation induced by BSA-FA@SP2 nanofluorophores. (a) Schematic representation of the experimental workflow for complement activation analysis. BSA-FA@SP2 nanofluorophores were incubated with plasma, followed by quantification of complement activation *via* a C3a-specific ELISA assay. (b) Simplified illustration of complement cascade activation, highlighting cleavage of C3 into C3a, where C3a acts as an anaphylatoxin associated with inflammatory responses. Thresholds for nanoparticle-induced complement activation are defined as baseline (stealth behavior), ∼2x baseline (partial activation), and >> baseline (immunogenic response). (c) Standard calibration curve for C3a ELISA showing absorbance at 450 nm as a function of C3a concentration (ng/mL), demonstrating strong linearity (R^2^ = 0.93). (d) Fold-change in C3a concentration following incubation of BSA-FA@SP2 nanofluorophores with plasma at varying concentrations (10% and 50% plasma). No statistically significant increase (ns) in C3a levels was observed compared to plasma-only controls, indicating minimal complement activation. Data are presented as mean ± standard deviation (n = 2).

Quantitative analysis of C3a generation revealed no statistically significant increase (ns) in complement activation for BSA-FA@SP2 nanofluorophores relative to plasma-only controls across both diluted (10%) and concentrated (50%) plasma conditions (**Figure 10d**). The observed fold-change remained approximately at baseline (∼1-1.3x), indicating minimal induction of complement activation. Based on established interpretation thresholds, nanoparticle-plasma interactions that remain at or near baseline levels are indicative of “stealth” behavior, whereas elevated signals (≥ 2x baseline) reflect partial activation and significantly higher signals denote immunogenic responses (**Figure 10b**). Accordingly, the BSA-FA@SP2 nanofluorophores demonstrates a non-immunogenic, complement-inert profile under the tested conditions.

These findings are significant because complement activation is a key determinant of nanoparticle fate, influencing C3 opsonization, phagocytic clearance, and infusion-related inflammatory responses. Prior nanoparticle studies show that complement proteins, particularly C3 fragments, deposit on nanoparticle coronas and promote immune recognition and clearance.^66^ In addition, synthetic “stealth” coatings such as PEG or PEG-like polymers can still activate complement, including C3 cleavage and C3a release, depending on surface chemistry and serum conditions.^67^ In contrast, the negligible C3a response observed here suggests that the albumin-chaperoned BSA-FA@SP2 architecture limits complement engagement.

Although direct reports specifically measuring C3a release from albumin nanocarriers remain limited, the broader literature supports this mechanism. Pre-formed albumin coronas have been shown to decrease complement activation and inhibit nonspecific plasma protein adsorption, supporting the role of albumin-rich interfaces as protective biomimetic coatings.^68^ Separately, nanocarrier surface chemistries that preserve adsorbed albumin structure can reduce macrophage clearance, whereas denatured albumin conformations promote recognition by scavenger receptors.^69^ Therefore, the low C3a generation observed here may arise from native-like BSA surface presentation and shielding of the SP2 core, which together reduce nonspecific protein adsorption and downstream complement amplification.

Collectively, these results demonstrate that BSA-FA@SP2 nanofluorophores exhibit excellent hemocompatibility with respect to complement activation, reinforcing their classification as a stealth nanofluorophores. This property is critical for systemic administration, as it supports prolonged circulation, reduced immune clearance, and enhanced tumor-targeting efficiency, key requirements for translational NIR-II image-guided surgical applications.

## 3. Conclusion

In this study, we successfully engineered a dual-receptor-targeted NIR-II nanofluorophore (BSA-FA@SP2) designed to overcome the precision limitations of current ovarian cancer intraoperative guidance. By utilizing a BSA scaffold as a molecular chaperone, we achieved significant NIR-II fluorescence enhancement and robust NIR-II stability in complex biological fluids, including whole blood. This chaperoned architecture not only preserved emission integrity but also mitigated nonspecific protein interactions, enabling reliable signal performance under physiologically relevant conditions.

Through a systematic multi-scale validation framework, spanning *in vitro* 2D cellular models, 3D multicellular spheroids for interstitial penetration, and 3D tumor-on-a-chip microfluidics for flow-dependent transport kinetics. This comprehensive approach confirms that BSA-FA@SP2 nanofluorophores maintains targeting integrity and extravasation capability under physiological shear stress, providing a robust, animal-free validation of its translational potential. Importantly, our results confirm that the dual-targeting of SPARC and FRα receptors significantly increases uptake compared to non-functionalized controls, while maintaining excellent hemocompatibility and negligible cytotoxicity.

Critically, the nanofluorophores exhibited excellent biocompatibility, characterized by negligible cytotoxicity, minimal hemolysis, and complement-inert behavior indicative of a “stealth” profile. These properties are essential for systemic administration, as they support prolonged circulation and reduced immune clearance while preserving targeting efficiency.

Collectively, this work establishes a robust, animal-free paradigm for the development and validation of next-generation optical nanoprobes. By integrating biomimetic design with advanced human-relevant testing platforms, this strategy provides a clear and scalable pathway toward clinically translatable, high-fidelity NIR-II image-guided surgical navigation for epithelial ovarian cancer and other complex malignancies.

## 4. Experimental Section/Methods

*Materials:* All materials were purchased from Sigma-Aldrich and used without modification, unless otherwise stated. Porcine tissues were obtained from local grocery stores. Preserved ovine porcine ovaries were purchased from Nasco Education.

*In Silico Investigation of Protein-Dye Interactions:* Molecular docking stimulations were performed to better understand binding affinity and molecular configuration between PCPDTBT and BSA. The monomer unit of PCPDTBT was generated using Avogadro and docked within the crystal structure of BSA (PDB ID: 4F5S) using UCSF Chimera and AutoDock Vina. Binding affinity (in kJ/mol) was collected using UCSF Chimera software, and molecular interactions were considered for all amino acids within 4.5 of the PCPDTBT monomer unit. The following grid dimensions were used for each globular protein analyzed in the Chimera software. HSA: grid center (27.495, 7.07482, 77.542) and size (91.3411, 21.806, 72.38). BSA: grid center (33.9247, 25.1865, 95.3462) and size (132.603, 58.6942, 84.3907).

LF: grid center (-7.52873, 26.318, 8.60424) and size (81.2127, 58.054, 62.9955). Hb: grid center (9.4075, 14.4314, 40.2025) and size (60.871, 50.2431, 55.7531). Tf: grid center (43.0147, 45.3121, 49.5634) and size (54.2345, 64.134, 45.3631). ß-LG: grid center (11.0884, 9.37318, 36.9725) and size (35.4169, 39.3696, 34.525).

*Fluorescence Titration Assay for BSA-PCPDTBT Binding:* To evaluate the binding affinity of PCPDTBT with BSA, PCPDTBT was first prepared at 0.5 mg/ mL in PBS1x at pH 7.4. This fixed PCPDTBT solution was incubated with increasing concentration of BSA (0-20 µM) at room temperature for 30 minutes to enable equilibrium binding. Each mixture was vortexed for 10s and shaken to ensure homogenous complexation. Fluorescence spectra was collected after incubation using the RF-6000 Spectro Fluorophotometer (Shimadzu) at λ_ex_: 645 nm, and the maximum emission intensity was recorded for each titration point. The binding curved was plotted as fluorescence intensity (a.u.) versus BSA concentration (µM). Data was fitted to a one-site binding hyperbola model using GraphPad Prism 9.0 to obtain the equilibrium dissociation constant (K_d_) and maximum binding capacity (*B*_max_). All experiments were performed in triplicate, and data is presented as mean ± SD.

*Preparation of DSPE-PEG@SP2:* To prepare DSPE-PEG@PCPDTBT, a previously established protocol reported by our group was used.^37^ 1 mg of PCPDTBT (Sigma-Aldrich) was codissolved in 2 mL of THF with 5 mg of DSPE-PEG2k (Biopharma PEG). This organic solution was added dropwise to an 8 mL PBS1x solution under probe sonication for 4 minutes. To evaporate the THF in solution, the solution stirred overnight at 450 rpm at room temperature overnight. The solution was then ultrafiltered through a 30 kDa filter at 4500 rpm and 4°C for 10 min three times with PBS1x washing to remove any excess precursors. The final product was collected after two successive filtrations through a 0.22 μm PVDF membrane and stored at 4 °C.

*Preparation of BSA@SP2:* To prepare BSA@PCPDTBT, a previously established protocol reported by our group was used.^37^ 1 mg of PCPDTBT was dissolved in 2 mL of THF. 52.8 mg of BSA (Sigma-Aldrich) was dissolved in 8 mL of PBS1x in a separate vial and stirred at 37°C until fully dissolved. The organic solution was added dropwise to the BSA solution under probe sonication for 4 minutes. To evaporate the THF in solution, the solution stirred overnight at 450 rpm at room temperature overnight. The solution was then ultrafiltered through a 100 kDa filter at 4500 rpm and 4°C for 10 min three times with PBS1x washing to remove any excess precursors. The final product was collected after two successive filtrations through a 0.22 μm PVDF membrane and stored at 4 °C.

*EDC/NHS Crosslinking of FA and BSA:* To functionalize the primary amines of BSA with folic acid (FA, Sigma-Aldrich), an EDC/NHS crosslinking method was used.^37^ 52.8 mg of BSA was dissolved in 8 mL of PBS1x and stirred at 37°C until fully dissolved. In a separate vial, 20 mg of FA was dissolved in 2 mL of dimethyl sulfoxide. The FA solution was then added to a vial with 43.4 mg of 1-Ethyl-3-(3-dimethylaminopropyl) carbodiimide hydrochloride (EDC·HCl, Millipore Sigma) and 26.1 mg of N-hydroxysuccinimide (NHS, Sigma-Aldrich) under constant stirring. 100 µL of triethylamine (TEA, Sigma-Aldrich) was added to the FA/EDC/NHS solution and allowed to stir for 20 minutes in the dark. The FA/EDC/NHS solution was then added dropwise to the BSA solution and measured at a pH ∼10. The final BSA-FA solution was stirred overnight in the dark and dialyzed in DI water for 24 hours using a 20k MWCO dialysis cassette.

*Preparation of BSA-FA@SP2:* To prepare BSA-FA@SP2, 1 mg of PCPDTBT was first dissolved in 2 mL of THF. This organic solution was added dropwise to the BSA-FA solution under probe sonication for 4 minutes. The solution was immediately ultrafiltered through a 100 kDa filter at 4500 rpm and 4°C for 10 min three times with PBS1x washing to remove any excess precursors. The final product was collected after two successive filtrations through a 0.22 μm PVDF membrane and stored at 4 °C.

*Physicochemical and Optical Characterization:* Hydrodynamic diameter and polydispersity index (PDI) were measured via dynamic light scattering (DLS), using the Litesizer DLS 700 Particle Analyzer instrument (Anton Paar). DLS measurements were recorded using 1:100 nanoparticle solution: PBS1x volume solutions. ζ-potential measurements also used the Litesizer DLS 700 Particle Analyzer. ζ-potential measurements were recorded using 1:4 nanoparticle solution: PBS1x volume solutions. Ultraviolet–visible-near-infrared (UV–vis-NIR) absorption was recorded using the Genesys 30 Visible Spectrophotometer (ThermoFisher Scientific). All absorbance measurements were recorded with scanning intervals of 1 nm from. Fluorescence spectra were collected on an RF-6000 Spectro Fluorophotometer (Shimadzu) at an interval of 1 nm scanning.

*Transmission Electron Microscopy Images:* Carbon coated copper grids of 200 mesh were glow discharged immediately prior to use to render the surface hydrophilic. A drop of PCA NP was then applied to the grid surface, and it was incubated for 60 seconds. Excess liquid was removed by blotting with filter paper. Negative staining was performed by adding a drop of freshly prepared 1% w/v uranyl acetate solution to the grid surface, followed by a 30-second incubation. Excess stain was then carefully blotted away, and the grids were dried under vacuum at room temperature overnight. All samples were imaged using a Hitachi H-7650 transmission electron microscope at 100 kV.

*Colloidal Stability:* The colloidal stability of BSA-FA@SP2 was evaluated by monitoring changes in hydrodynamic diameter and polydispersity index (PDI) over time using dynamic light scattering (DLS). Nanoparticle suspensions were prepared at a 1:100 dilution in PBS1x. Samples were stored under three conditions: 4°C (storage condition), 37°C (physiological temperature), and 37°C in the presence of plasma to simulate biologically relevant environments. DLS measurements were recorded at 24-hour intervals over a period of 7 days. Changes in particle size distribution and PDI were used as indicators of nanoparticle aggregation or degradation under each condition.

*Protein Quantification of BSA-FA@SP2 and BSA@SP2 Nanofluorophores via BCA Assay.* To protein content of BSA-FA@SP2 nanoparticles was quantified using a bicinchoninic acid (BCA) assay. A standard calibration curve was generated using albumin standards ranging from 0 – 2000 µg/ mL. Briefly, 25 µL of each standard and nanoparticle sample was added to separate wells in a 96-well plate (n = 3). Subsequently, 200 µL of the BCA working reagent was added to each well and incubated at 37°C for 15 minutes. Following incubation, the absorbance for each well was collected at 562 nm and their white light picture were taken.

*Protein Characterization of BSA-FA@SP2 Nanofluorophores by Sodium dodecyl-sulfate polyacrylamide gel electrophoresis (SDS-PAGE):* To confirm the presence of albumin component in BSA-FA@SP2 nanoparticles, an SDS-PAGE gel was used to visualize protein bands. A disulfide reducing agent was prepared by adding 2-mercaptoethanol (ßME) to 4X laemmli buffer at a 1:9 ratio. The ßME-Laemmli solution was added to each sample in a 1:3 ratio and centrifuged at 3000 rpm for ∼20 seconds. All samples were heated at 95°C for 8 minutes and centrifuged under the same conditions. 10 µL of ladder and 25 µL of sample was added to separate wells in a pre-made mini SDS-PAGE gel cassette. Gel electrophoresis was performed at 120 V for 90 minutes, and the gel was washed with DI water three times. The resulting gel was then imaged using the Bio-Rad ChemiDoc Go Imaging System.

*Cell Viability in 2D Ovarian Carcinoma Models:* To evaluate the effects of BSA-FA@SP2 on cellular metabolic activity, cell viability was assessed using a standard MTT assay. Human ovarian carcinoma cell lines were plated in a 96-well plate at 0.5 x 10^6^ cells/mL (OVCAR8, SKOV3, OVCAR3) and 0.5 x 10^5^ cells/mL (ID8) for 24h to reach approximately 80% confluency. Cells were incubated with BSA-FA@SP2 at concentrations ranging from 1 to 100 µg/mL. After 48 hours, 20 µL of a 5 mg/mL 3-(4,5-Dimethyl-2-thiazolyl)-2,5-diphenyl-2H-tetrazolium bromide (MTT, Millipore Sigma) solution was added to each well. After 4 hours of incubation, the media was replaced with 200 µL of DMSO to dissolve the formazan crystals and mixed for 5s before collecting absorption at 592 nm in a microplate spectrophotometer (Agilent BioTek). The following formula was used to calculate cell viability:

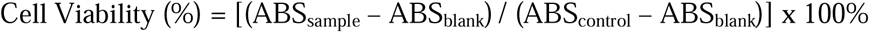

*Time-Dependent Cell Uptake in 2D Ovarian Carcinoma Models:* To evaluate time-dependent endocytosis of the experimental nanoprobes, human ovarian carcinoma cell lines (OVCAR8, SKOV3) were seeded in a 6-well plate at a density of 0.5 x 10^6^ cells/mL and cultured for 24h to reach approximately 80% confluency. Cells were treated with 100 µg/ mL of their respective nanoprobe treatment for 4, 24, and 48 hours. After their respective incubation time, the cells were washed with PBS1x and collected using trypsin 0.25%, 2.21 mM EDTA 1X. Cell suspensions were collected and imaged using an IVIS imaging system for all treatment groups at λ_ex_: 675nm and λ_em_: 840nm.

*Receptor Blocking Study in 2D Ovarian Carcinoma Models:* To evaluate receptor-mediated endocytosis of BSA-FA@SP2, human ovarian carcinoma cell lines (OVCAR8 and SKOV3) were seeded in a 6-well plate at 1 x 10^5^ cells/mL and cultured for 24 hours to reach approximately 80% confluency. Cells were then pre-treated with 10 µM of BSA (blocking SPARC receptors) or 10 µM of FA (blocking FR-α receptors) for 4h. Following receptor blocking, the culture medium was replaced with fresh medium containing 100 µg/mL BSA-FA@SP2 treatment. Cells cultured in high-glucose Dulbecco’s Modified Eagle Medium (DMEM, Fisher Scientific) without nanoparticle treatment served as controls. After 24h of incubation, the media was removed and the cells were washed with PBS1x to remove excess nanoparticles and subsequently detached using Trypsin 0.25%, 2.21 mM EDTA 1X. Cell suspensions were collected and imaged using an IVIS imaging system at λ_ex_: 675nm and λ_em_: 840nm.

*Microfluidic On-Chip System Ink Preparation and Fabrication:* Gelatin methacrylate (GelMA) was employed as a biocompatible extracellular matrix (ECM) analog for fabrication of the perfusable tumor constructs. The synthesis of GelMA was performed according to our previously reported protocol.^70^ For bioink preparation, lyophilized GelMA was dissolved in Dulbecco’s phosphate-buffered saline (DPBS) at 37 °C to obtain an 8% (w/v) solution. The photoinitiator lithium phenyl-2,4,6-trimethylbenzoylphosphinate (LAP; Sigma-Aldrich) was incorporated at a concentration of 0.3% (w/v) to enable photocrosslinking. To generate hollow, perfusable microchannels within the tumor model, Pluronic F-127 (poloxamer 407), a biocompatible thermoreversible polymer, was utilized as a sacrificial ink. A 40% (w/v) Pluronic F-127 solution was prepared by dissolving the polymer powder (Sigma-Aldrich) in DI water at 4°C to ensure complete solubilization. All prepared bioinks were stored at 4°C until further use in subsequent experiments.

A 3D printed polylactic acid (PLA) template (15 × 10 × 2 mm) was placed at the center of a 35 mm Petri dish to serve as a mold insert. Polydimethylsiloxane (PDMS; Sylgard 18) prepolymer mixture was then poured into the dish to fully encapsulate the PLA block. The assembly was cured at 70°C overnight in a conventional oven to allow complete crosslinking. The PLA template was then carefully removed, yielding a PDMS mold with a well-defined central cavity corresponding to the geometry of the printed insert. An 8% (w/v) GelMA solution was first cast into the PDMS mold to form a uniform bottom layer approximately 1 mm in thickness. Physical pre-gelation was induced by incubating the construct at 4°C for 10 minutes to promote rapid thermal gelation. Subsequently, a straight sacrificial filament (500 µm diameter) composed of Pluronic F-127 was printed within the mold using an extrusion-based 3D bioprinter (TissueStart, Tissuelabs), defining the future perfusable channel. A 5 µL aliquot of cell-laden bioink (1 × 10^6^ cells/mL) was then manually deposited adjacent to the printed sacrificial filament to localize the tumor cell population near the channel interface. An additional 1 mm thick layer of 8% GelMA was overlaid to encapsulate the construct, followed by ultraviolet (UV) crosslinking (10 mW/cm^2^ for 40 s) to stabilize the hydrogel network. The Pluronic F-127 sacrificial ink was subsequently liquefied at 4°C and carefully evacuated using a syringe, resulting in a hollow microchannel embedded within the tumor construct. Two 27 G needles were inserted at both ends of the channel and connected to a peristaltic pump via 0.5 mm inner-diameter tubing to enable continuous medium perfusion. The constructs were maintained in a humidified incubator at 37 °C, and culture medium was perfused at 16 µL/min, generating an estimated wall shear stress of ∼0.22 dyn/cm^2^, which falls within the physiologically relevant range reported for peritoneal tissues (0.02–1 dyn/cm^2^).^71^ Cells were allowed to proliferate and self-assemble into tumor clusters over a three-day period under perfusion. Thereafter, nanoparticles (100 µg/mL) were introduced into the perfusion medium and circulated through the channel for 36h. Following exposure, the constructs were perfused with fresh culture medium for 24h to remove residual nanoparticles from the channel and surrounding microenvironment.

*Tumor-On-Chip Cell Cluster Growth and Cell Viability:* The human ovarian carcinoma cell line SKOV3 (ATCC) was maintained and expanded following standard culture protocols as previously described.^72^ Prior to bioprinting, cells at appropriate confluency were detached using 0.25% (w/v) trypsin–EDTA, collected, and centrifuged to obtain a cell pellet. The cells were subsequently resuspended in 8% (w/v) GelMA prepolymer solution to prepare a cell-laden bioink at a final density of 1 × 10^6^ cells/mL for downstream fabrication of the tumor constructs.

Cell viability was assessed following 36h of nanoparticle exposure. A standard live/dead staining assay was performed in accordance with previously established protocols ^73^. Viability was quantified by calculating the percentage of live cells relative to the total cell population within each sample. Cell viability was quantified using a custom image analysis workflow implemented in MATLAB (MathWorks). Fluorescence images from live/dead staining were separated into green (live cells) and red (dead cells) channels. Each channel was processed using automated thresholding (Otsu’s method) to generate binary masks, followed by noise removal and hole filling to improve segmentation accuracy. Connected component analysis was then applied to identify and count individual cells in each channel. Cell viability was calculated as the percentage of live cells relative to the total number of cells (live + dead) for each image. Quantification results were visually validated by overlaying detected cell centroids onto the original images.

*3D Spheroid Ovarian Carcinoma Models:* OVCAR3 human high-grade serous ovarian cancer cells (ATCC, Manassas, VA) were grown in Roswell Park Memorial Institute (RPMI) 1640 supplemented with 10% heat-inactivated fetal bovine serum (FBS, Peak Serum, Inc., Wellington, CO), and 1X Antibiotic-Antimycotic solution. ID8 murine ovarian cancer cells (Cytion, Heidelberg, Germany) were grown in Dulbecco’s Modified Eagle Medium (DMEM) supplemented with 5% heat-inactivated FBS, 1X Insulin-Transferrin-Selenium (ITS), and 1X Antibiotic-Antimycotic solution. Cell lines were maintained at 37°C with 5% CO2, with routine sub-cultures. Cell lines were used between passages 1-25. Ovarian cancer spheroids were formed in a 384 well hanging drop array based on previously established protocols.^74–79^ Briefly, ovarian cancer spheroids were generated with OVCAR3 or ID8 cells with 1000 cells/drop in hanging drop arrays and maintained for up to 6 days. Phase contrast microscopy was used to monitor spheroid formation, compaction, and growth over 6 days.

*Cell Viability in 3D Ovarian Carcinoma Spheroids:* Nanoparticle doses of 0, 10, and 25 μg/mL were added to spheroids by diluting nanoparticle stock solutions 1:4 in the spheroid droplet. The CellTiter Glo Luminescent Viability ATP reporter assay (Promega Corporation, Madison, WI) was used to assess spheroid viability following 24 hours or 48 hours of nanoparticle exposure. Individual spheroids were transferred to a 384-well opaque-walled plate for luminescence readings. The CellTiter Glo assay was performed according to manufacturer’s protocols, using a 1:1 volume of reagent to spheroids. Plates were mixed for 2 minutes to lyse the cells with the CellTiter Glo reagent and incubated for 10 minutes at room temperature, prior to reading luminescence on a BioTek Cytation 7 microplate reader. Statistical analysis was performed with GraphPad Prism 10 software. All reported values represent the mean and standard error of the mean taken from 3-6 biological replicates, with at least three technical replicates within each experiment. Spheroid percent viability was determined by normalizing luminescence values to the average 0 μg/mL nanoparticle-treated spheroid luminescence and multiplying by 100%.

*Time- and Concentration-Dependent Cell Uptake in 3D Ovarian Carcinoma Spheroids:* Following 24 hours or 48 hours of nanoparticle exposure, 20 spheroids were pooled and spun down at 1000 x g for 5 minutes. Excess nanoparticle-dosed growth medium was removed manually by pipetting, and the spheroid pellet was rinsed and re-centrifuged in PBS1x to wash any residual soluble nanoparticles. Spheroid pellets were resuspended in radio immunoprecipitation assay (RIPA) buffer with 1X HALT protease cocktail (RRID:SCR_014485) to prevent protein degradation and maintained on ice for 1 hour with periodic vortexing to promote spheroid lysis. To gather nanoparticle fluorescence measurements, lysates were distributed in clear bottom, opaque walled 96 well plates. Plates were imaged using an IVIS imaging system at λ_ex_: 675nm and λ_em_: 840nm. Data shown in Figure S15 was imaged using a LI-COR Odyssey M imager at λ_ex_: 785 nm and λ_em_: 816-840 nm, and fluorescence was quantified with Image Studio software (version 6.1.0.79, LICORbio).

*Optical Performance in Biological Mileu:* To evaluate potential fluorescence quenching and stability of BSA-FA@SP2 in biological relevant environment, NP suspensions (100 µg/ mL) were incubated with various biological media, including bovine plasma (Sigma-Aldrich), 10% (v/v) packed porcine red blood cells (Innovative Research), and 10% (v/v) fetal bovine serum (Corning™). All samples were prepared in microcentrifuge tubes and maintained at room temperature during incubation. Following incubation, NIR-II fluorescence emission was collected at λ_ex_: 808 nm and λ_em_: 1250 – 1600 nm using the IR VIVO pre-clinical imager (Photon Etc.).

*Simulating Epithelial Ovarian Cancer with 3D Bioprinted Tumor-Mimicking Phantoms:* Epithelial tumor phantoms were 3D bio-printed in order to simulate epithelial ovarian cancer tumors. They were printed using our labs tumor-mimicking phantom solution which consists of intralipid (90 µL), hemoglobin (150 mg), gelatin (720 mg), and 17 ml PBS 1X. In which the hemoglobin acts as the light absorption component and intralipid acts as the light scattering component. By increasing and decreasing the concentration of intralipid (1x, and 2x) this will allow for closer optical mimicking of high and low grade epithelial ovarian tumors. The multiple solutions are then added to a syringe and added into the Bio One 3D Printer (Cellink). By inputting the following printing parameters: 10 mm/s speed, 25 µL pre-flow and extra pre-flow volumes, 100% infill extrusion multiplier, 3s post flow stop time, 10 µL/s extrusion rate, 25 µL retraction volume, 60 µL/s retraction rate, 3 mm Z-lift, 0.2s Z-offset, and 80% density, the ‘epithelial’ shape of the ovarian tumors can be replicated in our tumor mimicking phantom. Printing was done using cube dimensions 1 mm x 1 mm x 0.1 mm.

*Ex Vivo Porcine Ovary Simulating NIR-II Imaging of Epithelial Ovarian Cancer:* Fresh porcine ovaries (NASCO Education) were used as an *ex vivo* model to simulate epithelial ovarian tumor visualization. To mimic the superficial localization of epithelial tumors, 100µL of BSA-FA@SP2 (at 100 µg/ mL) was administered *via* microinjection into the outer cortical region of the ovary at an approximate depth of 1-2 mm from the surface. Care was taken to distribute the injection within a shallow, localized region to approximate tumor-like fluorescent domain rather than a single focal bolus. Following probe administration, whole-ovary NIR-II fluorescence imaging was performed using the IR VIVO preclinical imaging system with λ_ex_: 808 nm and λ_em_: 1250 – 1600 nm. To evaluate contrast between the simulated tumor region and surrounding normal ovarian tissue, fluorescence intensity profiles were analyzed using ImageJ. A linear region of interest (ROI) was drawn across the injected region and adjacent non-injected tissue. The NIR-II fluorescent radiant intensity along the line was extracted and plotted as a function of distance to assess signal distribution, contrast, and spatial delineation between the simulated epithelial tumor and healthy tissues.

*Ex Vivo Tissue Penetration in Tumor-Mimicking Phantoms:* To simulate an image-guided scenario, the optical environment of a tumor was simulated using a previously established protocol for spherically molded tumor phantoms containing DSPE-PEG@ and BSA@SP2.^61^ Porcine tissues slices at a thickness of 2 mm were used to simulate three distinct tissue types: skin (pork belly), muscle (pork tenderloin), and fat (pork chop) which each resembled the texture and composition of their respective tissue type (n=3). Tissues were stacked on a 12-well plate containing the tumor-mimicking phantoms for fluorescence imaging. IVIS images were collected at λ_ex_: 675 nm and λ_em_: 840 nm. Tumor phantom-to-background ratios were plotted using radiant efficiency (a.u.) values measured with the Living Image^TM^ software (Revvity).

*Hemocompatibility:* The hemolytic activity of BSA-FA@SP2 was evaluated using porcine red blood cells (RBCs). Nanoparticles suspensions were prepared at concentrations of 1, 5, 10, 50, and 100 µg/ mL in PBS1x. DI water and PBS1x served as positive (100% hemolysis) and negative (0% hemolysis) controls, respectively. Porcine packed RBCs were diluted to 10% w/v in PBS1x. Subsequently, 50 µL of the RBC suspension was added to 1 mL of each treatment or control group (n = 3 technical replicates per condition) and incubated for 1 hour at 37°C. Following incubation, samples were centrifuged at 3000 rpm for 20 minutes to pellet intact RBCs. The supernatants were collected, and absorbance was collected at 540 nm using spectrometer to quantify hemoglobin release. To further assess RBC membrane integrity, the cell pellets were resuspended in 1 mL of PBS1x and transferred to a clear, 12-well plate for visualization under 200x magnification using an inverted microscope (Laxco™) to qualitatively evaluate morphological changes. Hemolysis percentages were calculated using the following formula:

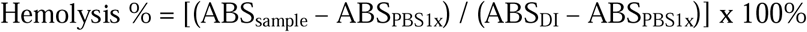

*Complement C3a Activation Assay.* Complement activation induced by BSA-FA@SP2 nanofluorophores was evaluated by quantifying the generation of C3a using a commercially available human C3a enzyme-linked immunosorbent assay (ELISA) kit (Thermo Fisher Scientific). The assay is based on a sandwich immunoassay format, in which C3a present in the sample is captured by immobilized anti-human C3a antibodies and detected via a biotin-conjugated secondary antibody followed by streptavidin-HRP and a colorimetric substrate, with absorbance measured at 450 nm. Plasma samples were prepared under cold conditions to minimize spontaneous complement activation and diluted to working concentrations of 10% and 50% (v/v) in assay buffer. BSA-FA@SP2 nanofluorophores were incubated with plasma for 30 minutes at 37 °C. Prior to ELISA, plasma samples were pre-diluted according to manufacturer recommendations (adjusted appropriately for plasma) to ensure measurements fell within the dynamic range of the assay. A seven-point standard curve was generated using serial dilutions of recombinant human C3a standard (0-20 ng/mL), and sample concentrations were interpolated using linear regression. Absorbance values were measured at 450 nm using a microplate reader, and C3a concentrations were normalized to plasma-only controls to determine fold-change in complement activation. All samples were analyzed in duplicate, and data are reported as mean ± standard deviation. Statistical significance was assessed using one-way ANOVA with post hoc comparisons, where p > 0.05 was considered not significant (ns). To interpret complement activation, fold-change values near baseline were classified as stealth behavior, approximately two-fold increases as partial activation, and substantially elevated responses as immunogenic, consistent with established nanoparticle–complement interaction frameworks.

## Supporting Information

Supporting Information is available from the Wiley Online Library or from the author.

## Author Contributions

I. S. conceived the idea and design the experiments. I.V. led the overall study. I. V. and I. S. analyzed all the data for this study except 3D spheroids. I.V., M.M., and J. C-X. were involved in data collection and curation, formal analysis, investigation, methodology, validation, and visualization of the experiments under the supervision of I.S. 3D bioprinting of TMPs were conducted by I. D. L. under the supervision of I.S. L.L.N. conducted all the 3D spheroids biocompatibility and cellular uptake measurements and quantification under the supervision of S.A.R. M.S conducted all the on-chip fabrication, biocompatibility studies, and cellular transport experiments under the supervision of C.X. The first draft of the manuscript was written by I.V. and I.S. with edits from all the authors. All authors have approved the final version of the manuscript.

## Acknowledgements

We would like to acknowledge funding for this work from the Edward E. Whitacre Jr. College of Engineering, Texas Tech University, and the American Heart Association (26CDA1597763). NIR-II imaging studies were conducted on the IR VIVO system, funded by the Cancer Prevention and Research Institute of Texas (CPRIT) at the Texas Tech University Health Sciences Center at Amarillo (Core Facility Support Award, RP200572). Spheroid cytotoxicity and uptake experiments were supported by the Gulf Coast Consortium Microphysiological Lead Optimization and Toxicity Screening facility (MLOTS RRID: SCR_023717) via CPRIT RP210108 (SAR). I.V. acknowledges the Barry Goldwater Scholarship and Excellence in Education Foundation. L.L.N acknowledges the support of the Texas A&M College of Engineering Horizons: 21 Fellowships for 21st Century Scholar doctoral fellowship. The authors thank Md. Hasnat Rashid for acquiring TEM images, and Nathaniel Bendele and Jonathan Djuanda for collecting tissue stacking IVIS images.

## Data Availability Statement

All the data associated with this work is either present in the main manuscript or the associated supporting information.

