## Supporting Information for "Modular Albumin-Chaperoned NIR-II Nanofluorophores Enables Pan-Ovarian Cancer Imaging Across Multiscale Tumor Models"

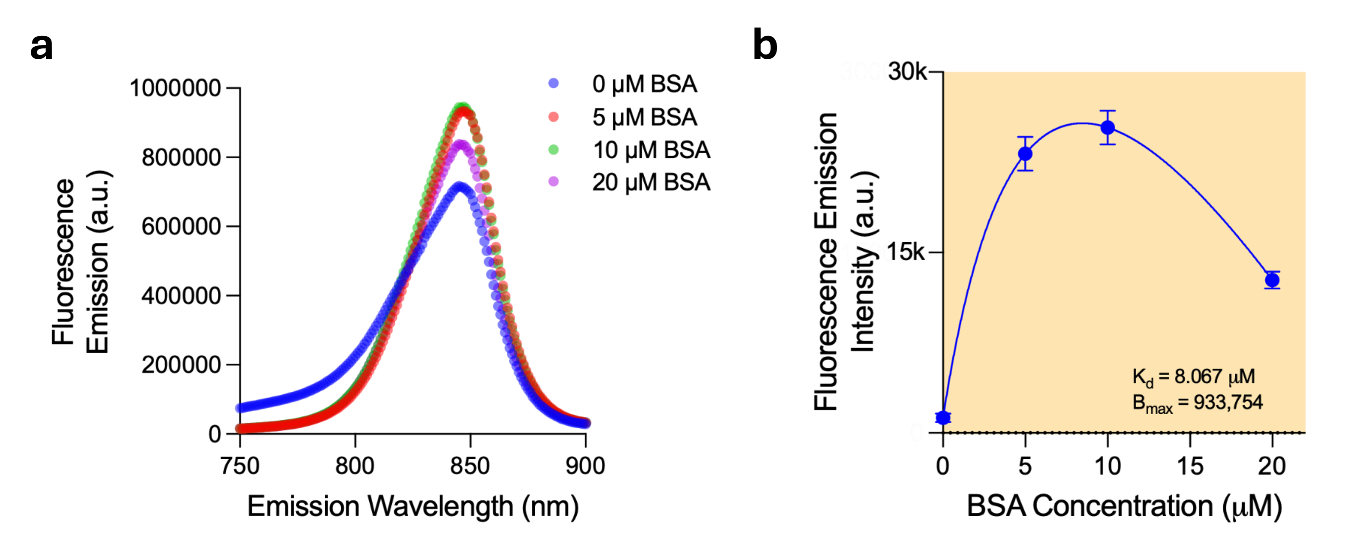

**Figure S1.** **Titration assay of PCPDTBT and BSA.** (a) Fluorescence spectra of PCPDTBT with increasing concentrations of BSA (0 – 20 µM). (b) Binding curve of PCPDTBT-BSA complex with a K_d_ of 8.067 µM and a B_max_ at 933,754.

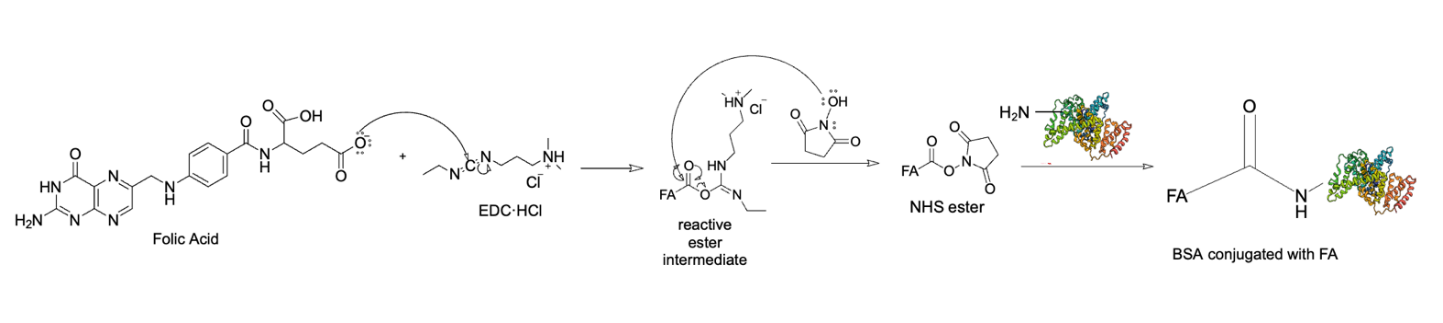

**Figure S2.** EDC/NHS crosslinking reaction scheme for BSA-FA conjugation.

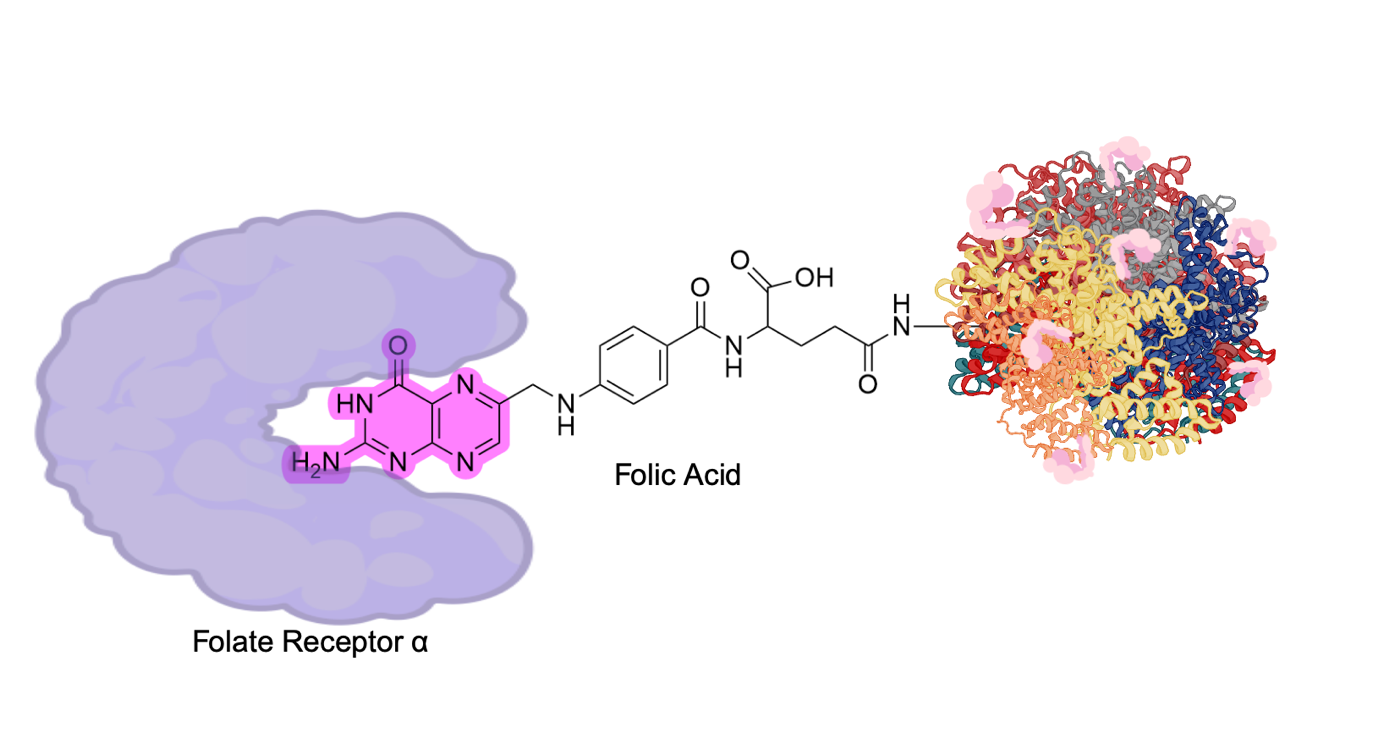

**Figure S3.** Binding orientation of FRα and BSA-FA, highlighting the interaction between folic acid’s pteridine ring and the FRα binding pocket.

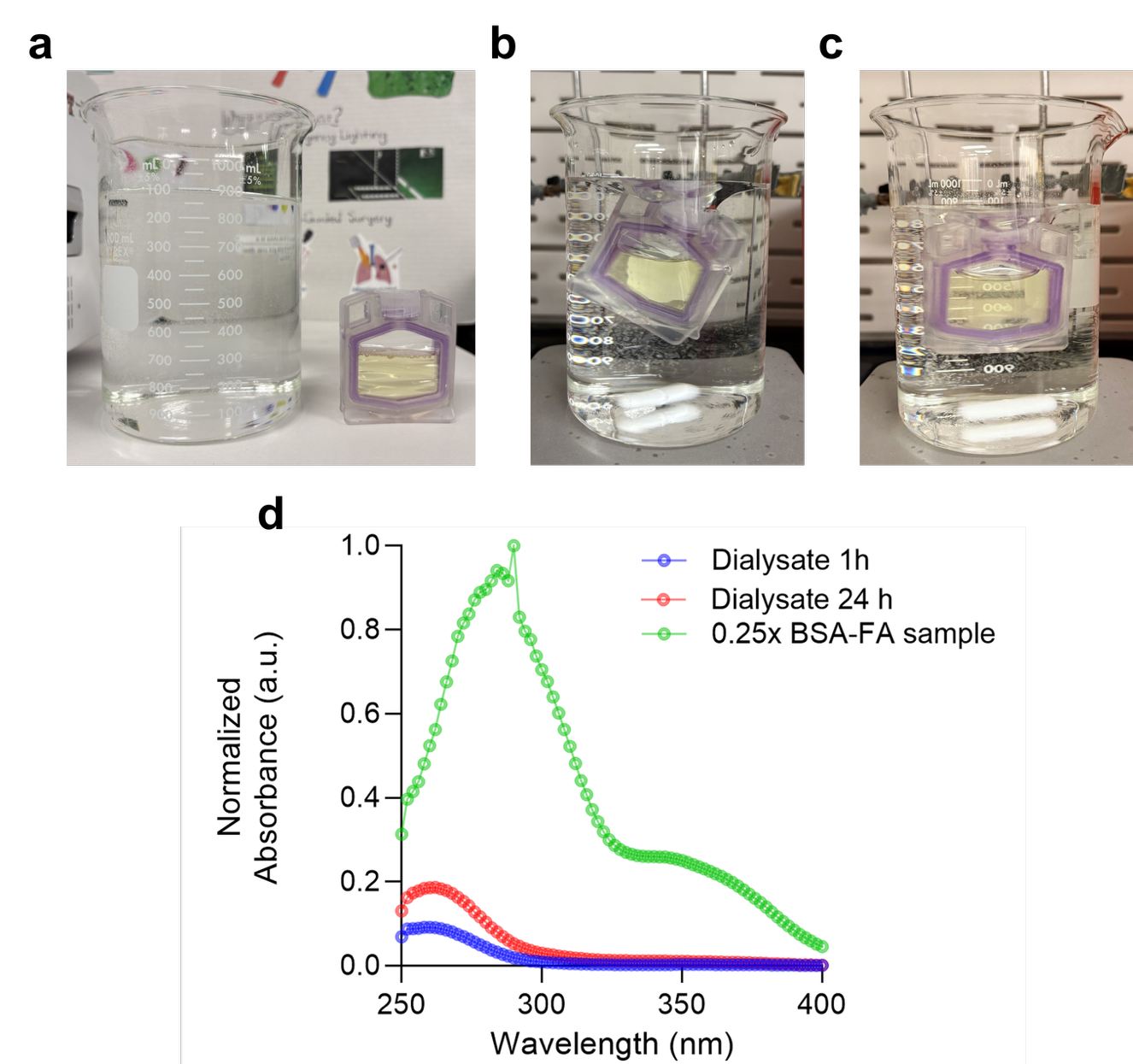

**Figure S4. BSA-FA purification *via* dialysis.** (a) BSA-FA solution in dialysis cassette and beaker containing dialysis solvent (DI water). Dialysis mixture at (b) 1 hour and (c) 24 hours with corresponding (d) UV-Vis absorbance spectra containing a BSA-FA character peak ~350 nm.

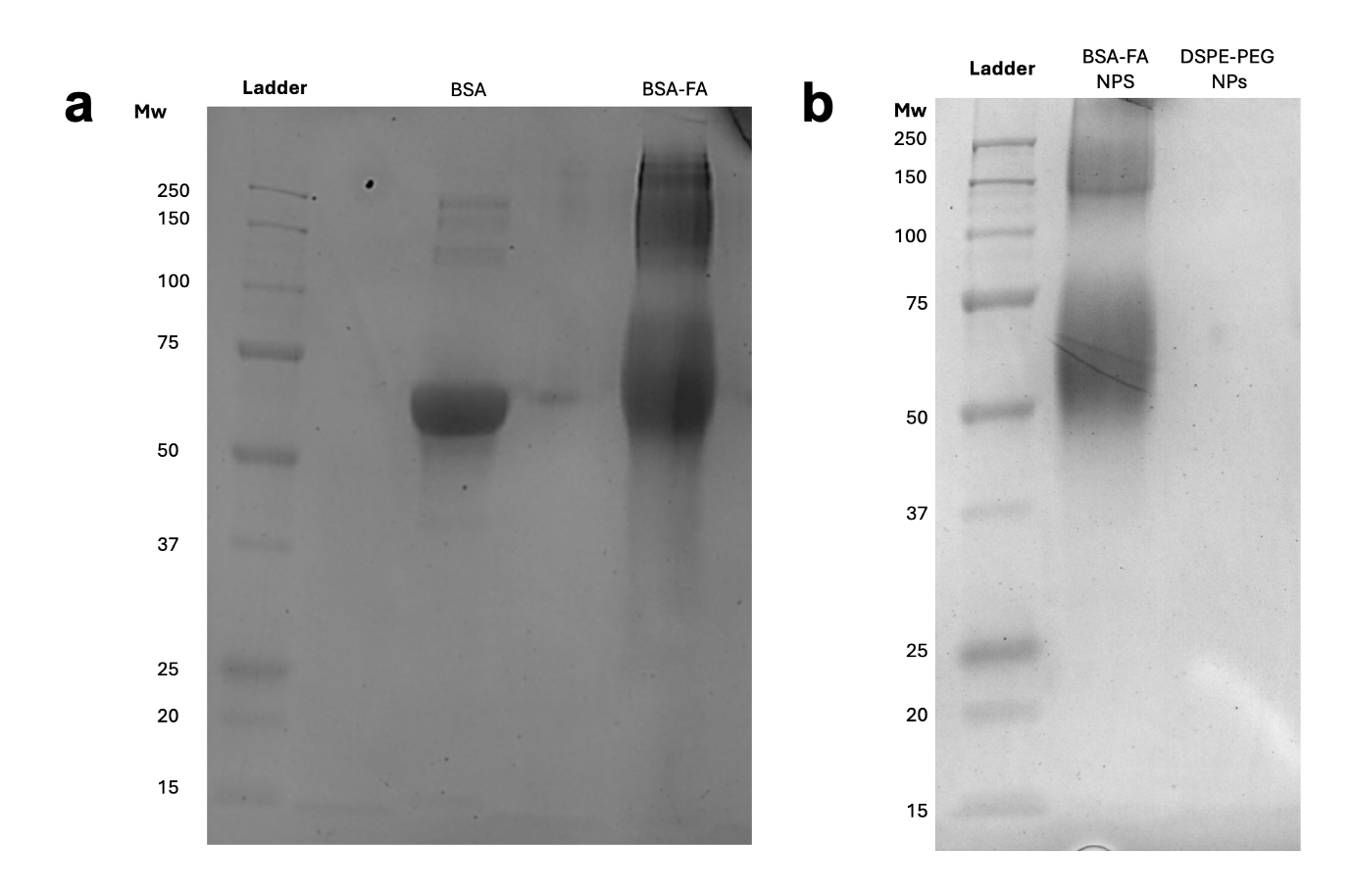

**Figure S5. Sodium dodecyl sulfate polyacrylamide gel electrophoresis (SDS-PAGE) of NPs.** SDS-PAGE results of (a) BSA and BSA-FA and (b) BSA-FA@SP2 and DSPE-PEG@SP2 with protein separation bands at ~66 kDa for BSA-based samples.

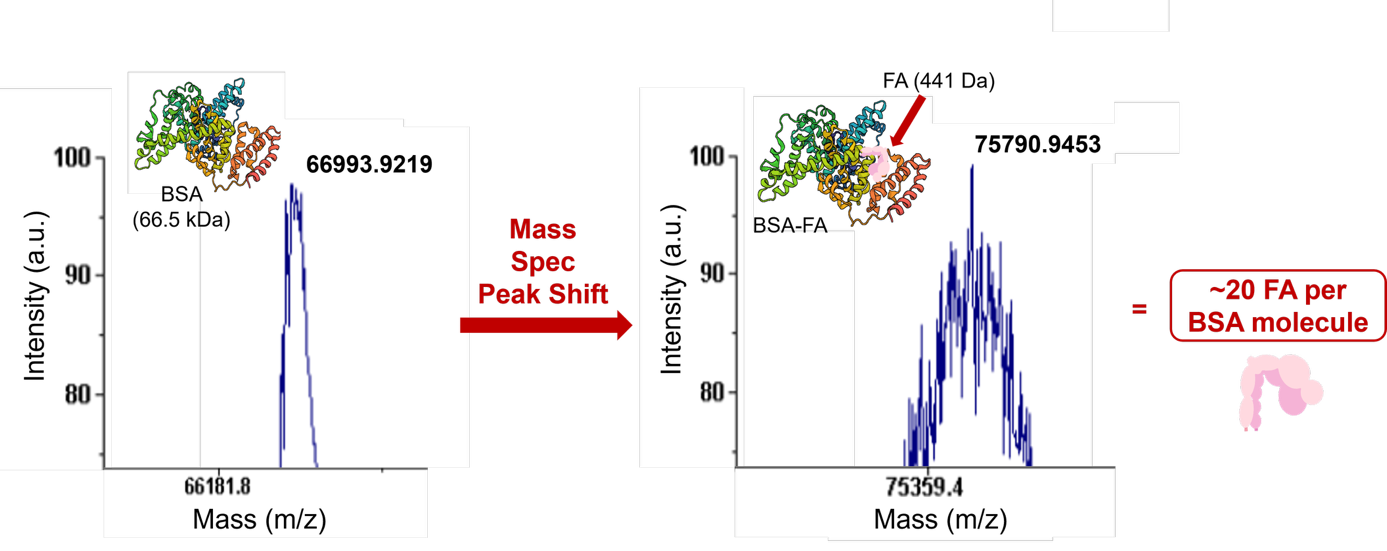

**Figure S6. Matrix-assisted laser desorption/ionization (MALDI-TOF)** mass spectra of native BSA and BSA-FA. The mass shift observed in the BSA-FA conjugate corresponds to an average density of ~20 FA molecules per BSA monomer.

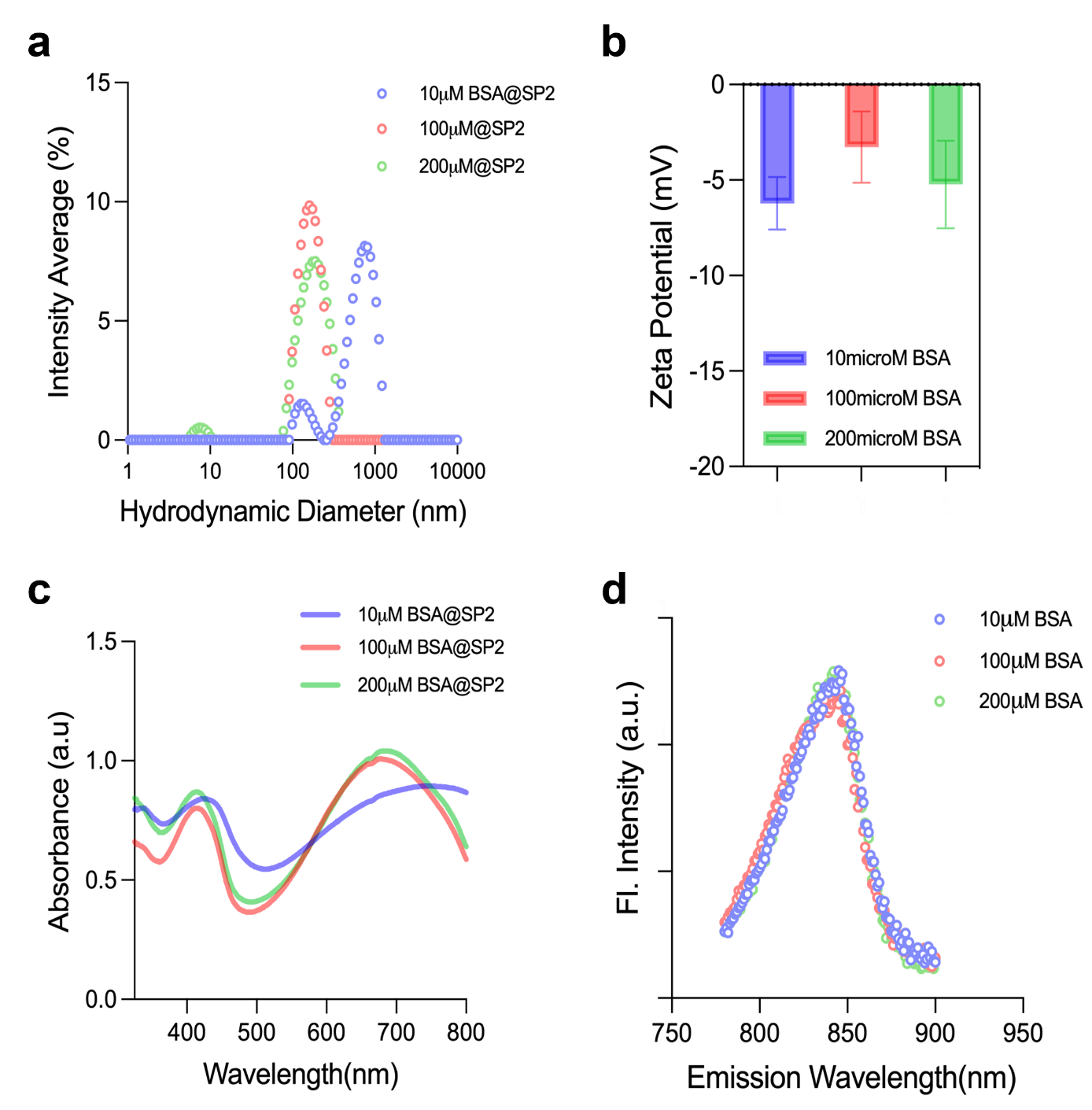

**Figure S7. Physicochemical and optical characterization of BSA NPs at increasing albumin concentrations.** (a) Hydrodynamic diameter measurements via dynamic light scattering, (b) zeta potential (in mV), (c) UV-Vis-NIR absorbance spectra, and (d) fluorescence emission spectra (λ_ex_ =645 nm) of 10µm, 100µm, and 200µm BSA coated SP2 nanoparticle solutions with fixed SP2 dye concentration.

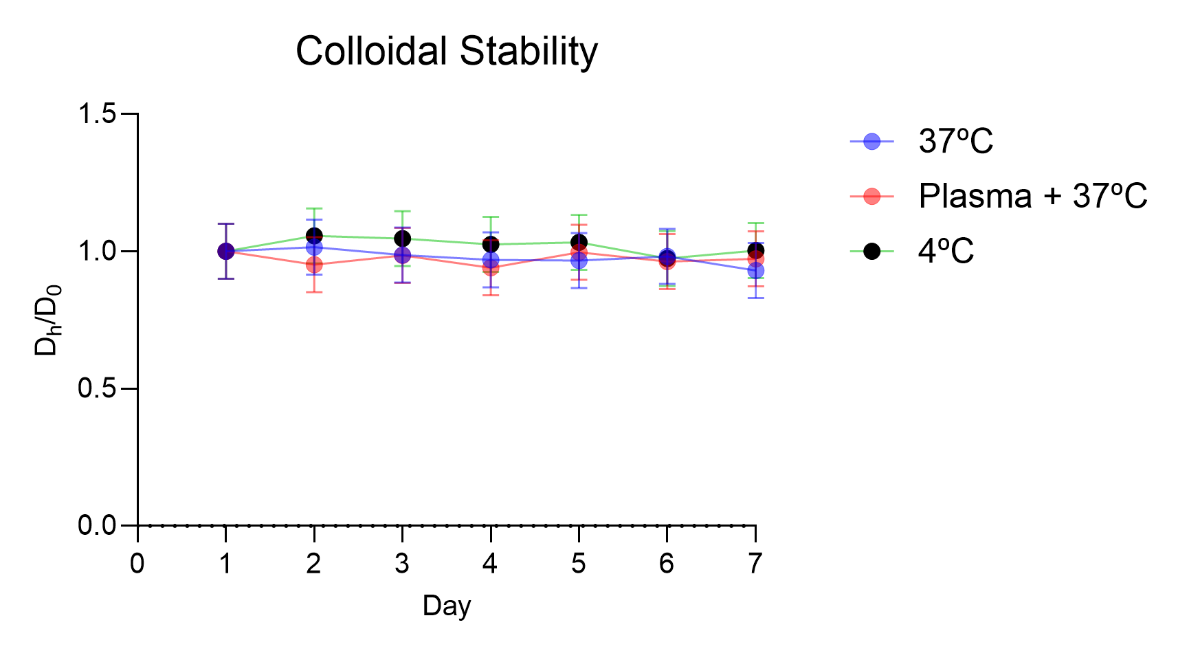

**Figure S8. Stability assessments of BSA-FA@SP2 nanofluorophores.** The nanofluorophores maintain a stable hydrodynamic diameter across physiological and storage conditions (4ºC, 37ºC, and plasma), indicating minimal aggregation.

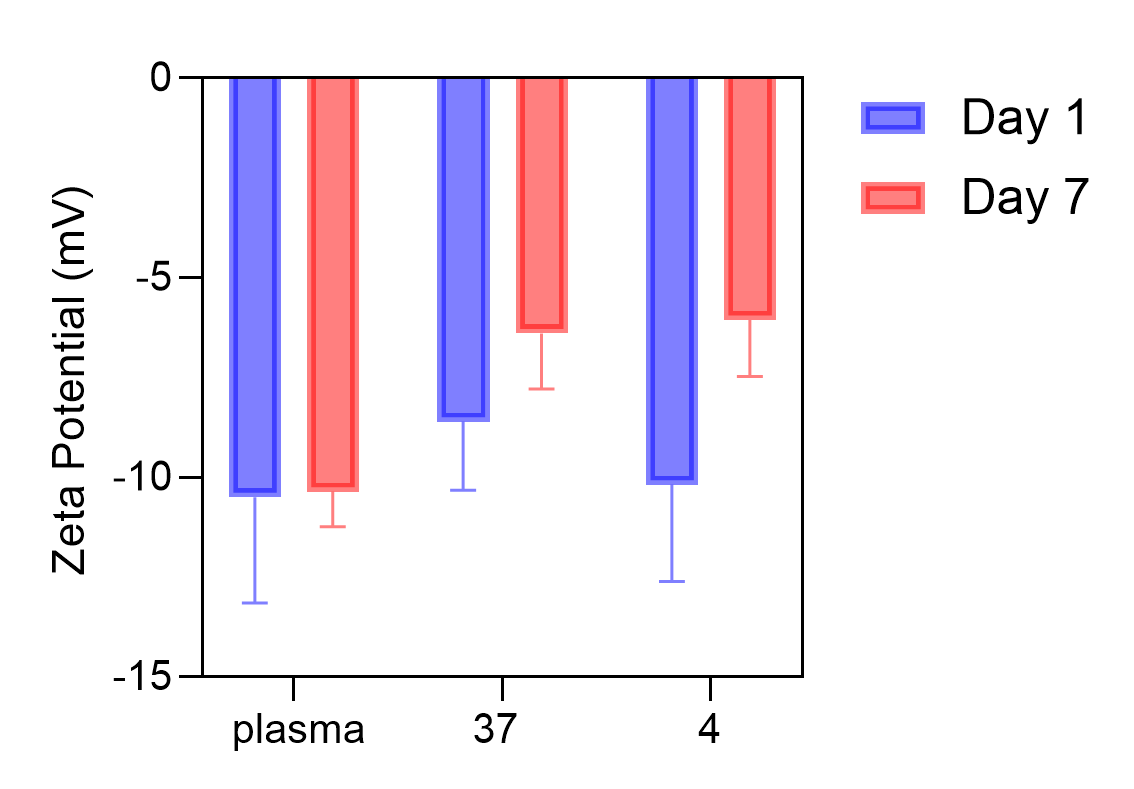

**Figure S9. Zeta potential stability of BSA-FA@SP2 nanofluorophores.** The surface charge was monitored under various storage and physiological conditions (4ºC, 37ºC, and plasma). Consistent negative zeta potential values across these environments indicate robust electrokinetic stability and resistance to significant protein-induced aggregation.

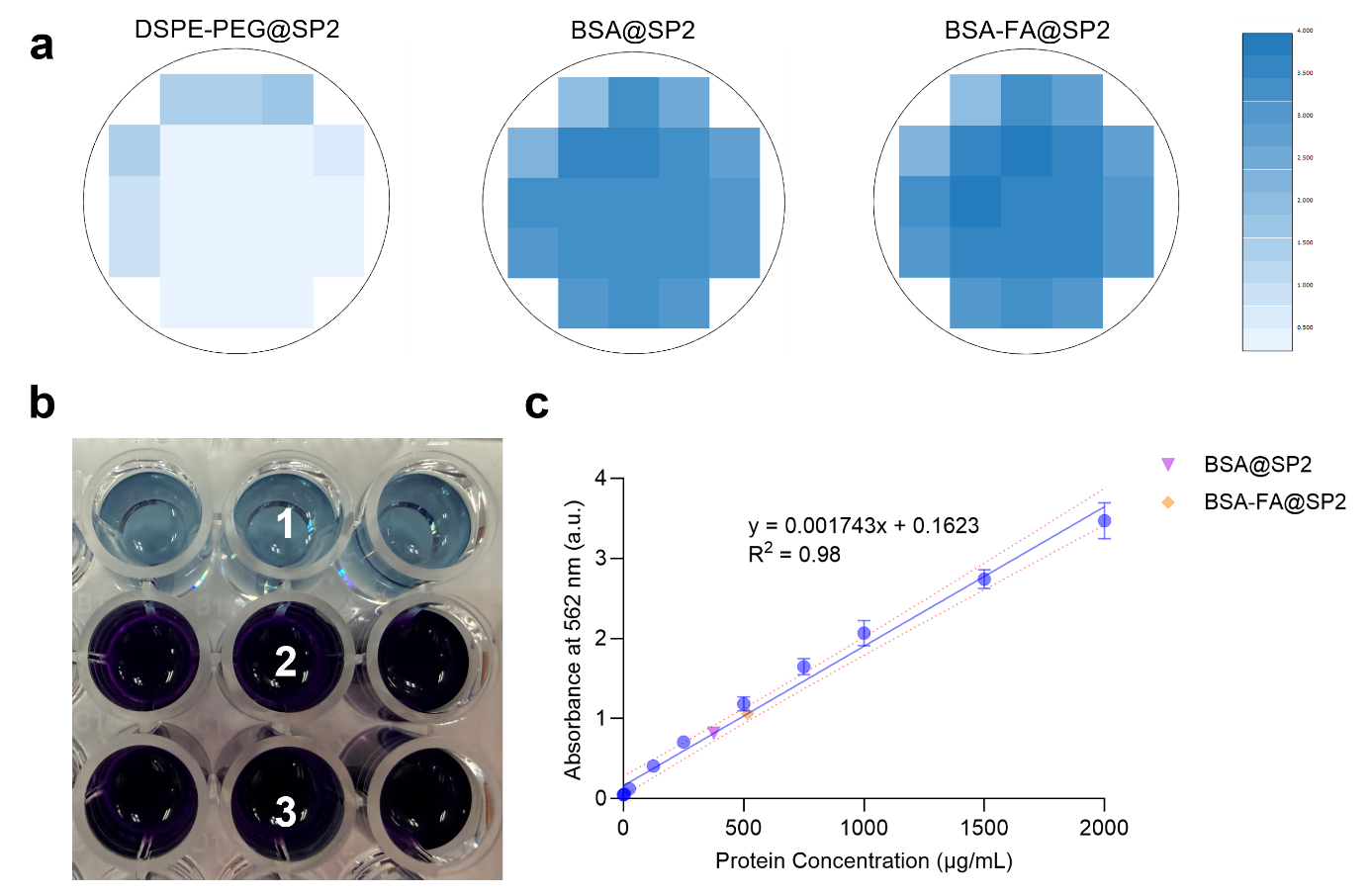

**Figure S10. Bicinchoninic acid (BCA) assay of nanofluorophore designs.** (a) Representative area scan of absorbance at 562 nm and (b) white light image of (1) DSPE-PEG@, (2) BSA@, and (3) BSA-FA@SP2 wells n = 3. (c) Protein standard curve with detectable levels of BSA in BSA@ and BSA-FA@SP2.

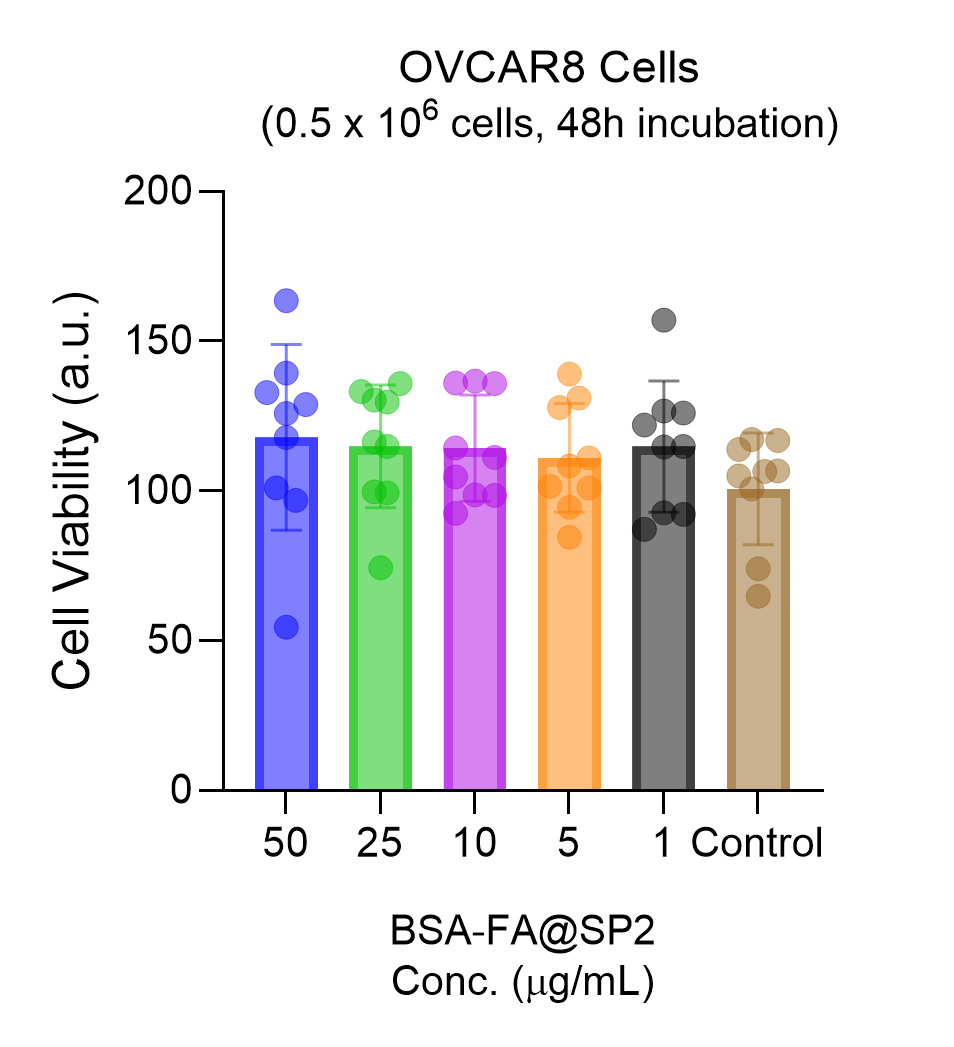

**Figure S11.** Cytotoxicity of BSA-FA@SP2 in 2D OVCAR8 model at concentrations 1 to 50 µg/mL, n = 9.

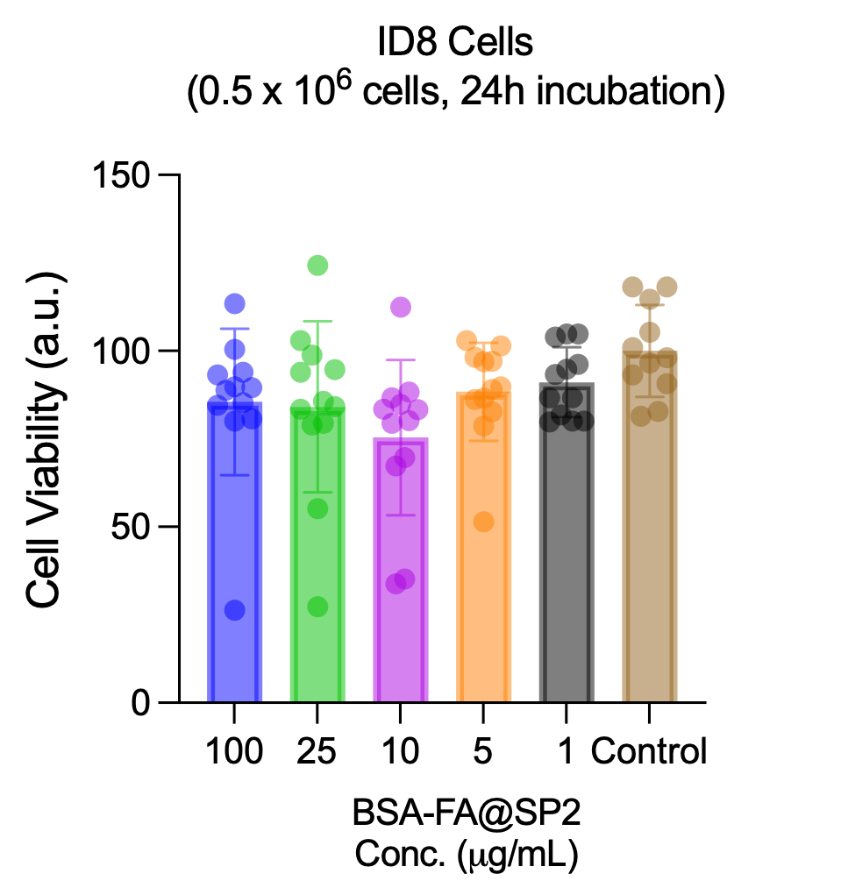

**Figure S12.** Cytotoxicity of BSA-FA@SP2 in 2D ID8 model at concentrations 1 to 100 µg/mL, n = 12.

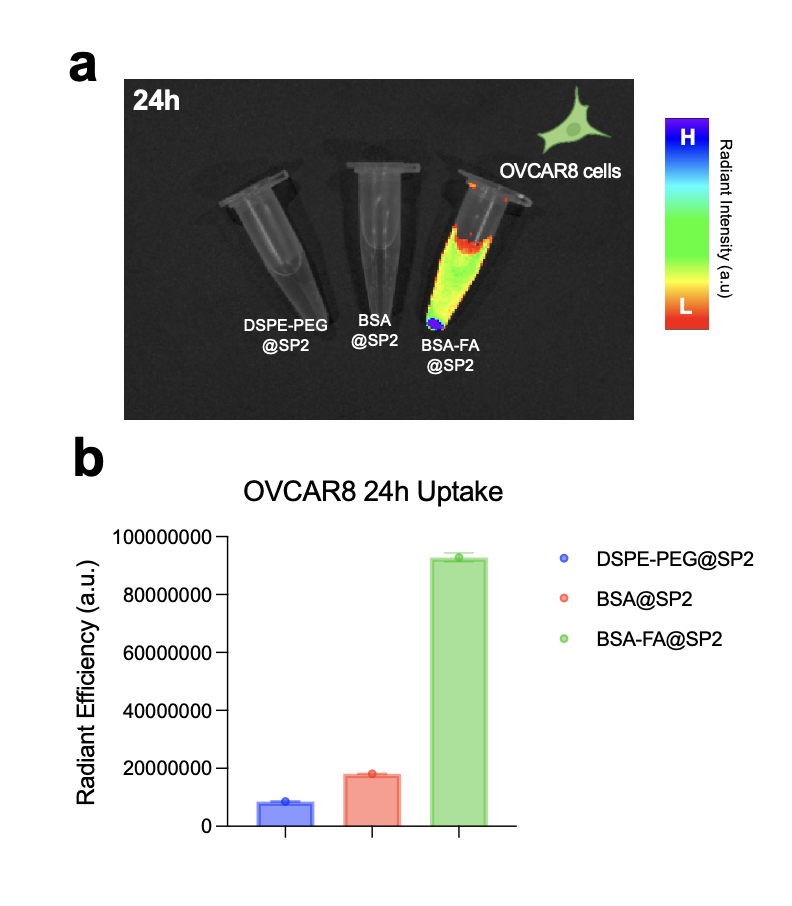

**Figure S13.** **24-hour uptake of nanofluorophores in OVCAR8.** (a) IVIS fluorescence images of OVCAR8 cultures after 24h treatment with DSPE-PEG@, BSA@, and BSA-FA@SP2 nanofluorophores, with (b) corresponding radiant intensities.

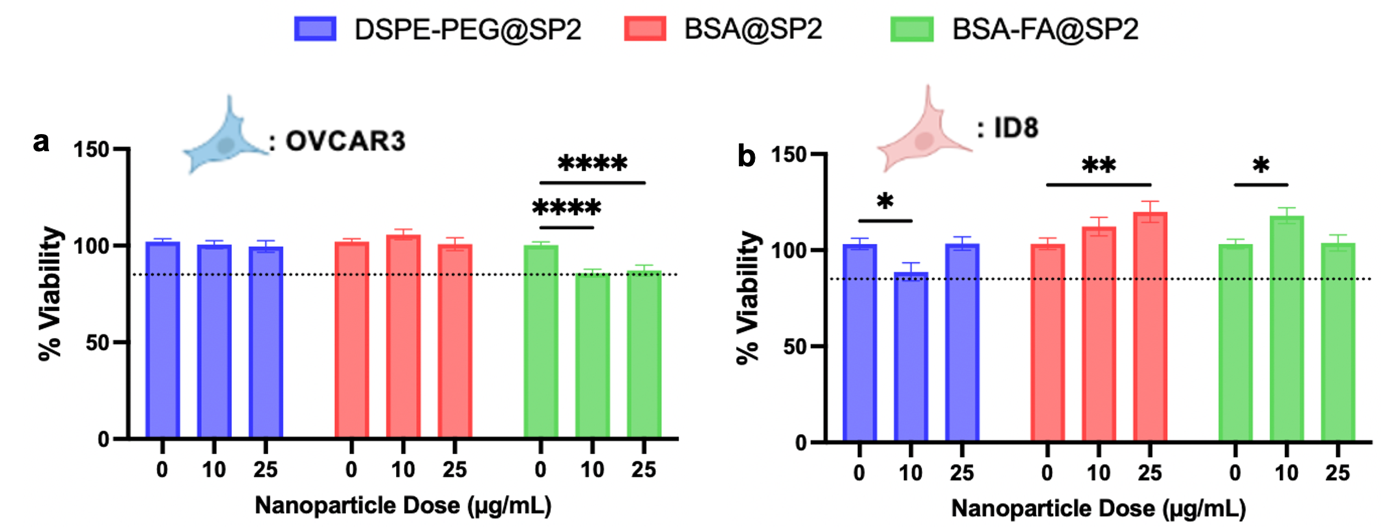

**Figure S14.** **3D spheroid viability after 48 hours of nanofluorophore exposure.** Dotted lines = 85% viability indicating a conventional cytocompatibility threshold. (a) OVCAR3 spheroid viabilities after culture with 0, 10, and 25 µg/mL of DSPE-PEG@SP2, BSA@SP2, and BSA-FA@SP2. All treatment conditions remained above the 85% viability threshold of acceptable cytocompatibility, with only minor decreases in BSA-FA@SP2 viability observed (****p<0.0001, two-way ANOVA). (b) ID8 spheroid viabilities after culture with 0, 10, and 25 µg/mL of DSPE-PEG@SP2, BSA@SP2, and BSA-FA@SP2. ID8 spheroids remained over 85% viable up to 48 hours after nanoparticle treatment, demonstrating extended cytocompatibility of each nanoparticle formulation, despite mild decreases in viability from untreated controls (*p<0.05, two-way ANOVA).

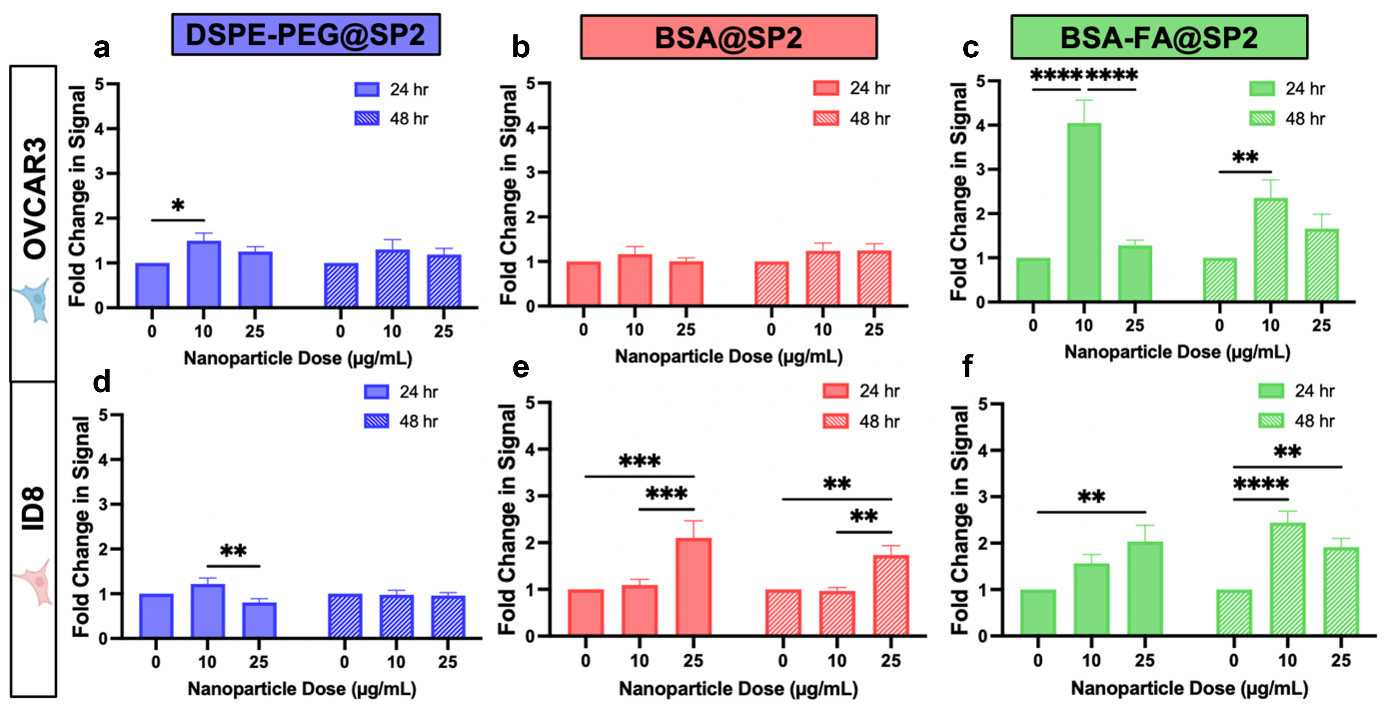

**Figure S15.** **24- and 48-hour uptake of nanofluorophores in 3D spheroids.** Graphs report fold change in fluorescent signal compared to an untreated control. Fold change in OVCAR3 spheroid lysate fluorescence after culture with 0, 10, and 25 µg/mL of a) DSPE-PEG@SP2 (blue), b) BSA@SP2 (red), and c) BSA-FA@SP2 (green). BSA-FA@SP2 had the highest uptake in OVCAR3 spheroids after both 24 and 48 hours (****p<0.0001, **p<0.01, two-way ANOVA). Fold change in ID8 spheroid lysate fluorescence after culture with 0, 10, and 25 µg/mL of d) DSPE-PEG@SP2 (blue), e) BSA@SP2 (red), and f) BSA-FA@SP2 (green). BSA-FA@SP2 had a maximum fold change of 2.04 after 24 hours and 2.44 after 48 hours of culture, demonstrating superior uptake compared to BSA@SP2 and DSPE-PEG@SP2 (**p<0.01, ****p<0.0001, two-way ANOVA).

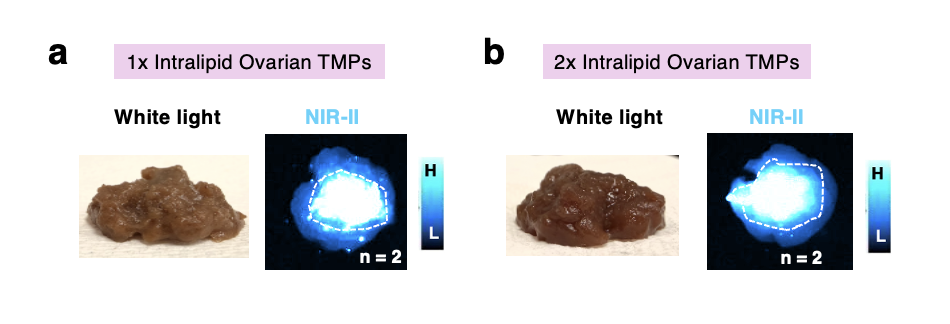

**Figure S16.** **3D-bioprinted tumor phantoms with BSA-FA@SP2, mimicking epithelial ovarian cancer.** White light (left) and NIR-II images (right) of tumor phantoms (a) 1X and (b) 2X Intralipid content.

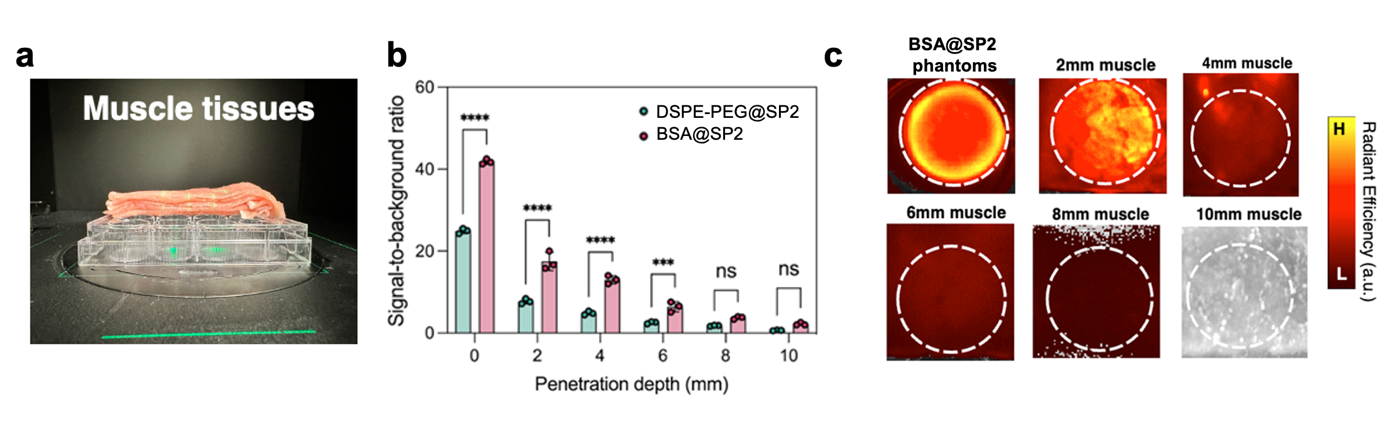

**Figure S17. Optical penetration of BSA@SP2 tumor-mimicking phantoms through porcine muscle.** (a) White light image of porcine muscle slices above tumor mimicking phantoms in a 12-well plate. (b) Fluorescence signal-to-background ratio and (c) IVIS images of tissue-covered tumor mimicking phantoms (n=3). IVIS images were collected at λ_ex_: 675 nm and λ_em_: 840 nm.

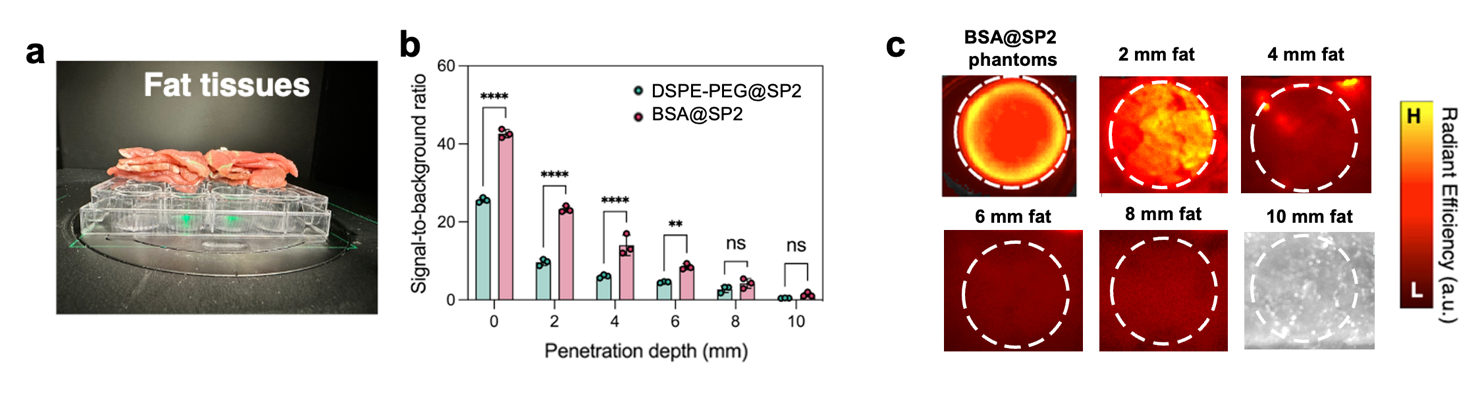

**Figure S18. Optical penetration of BSA@SP2 tumor-mimicking phantoms through porcine fat.** (a) White light image of porcine fat slices above tumor mimicking phantoms in a 12-well plate. (b) Fluorescence signal-to-background ratio and (c) IVIS images of tissue-covered tumor mimicking phantoms (n=3). IVIS images were collected at λ_ex_: 675 nm and λ_em_: 840 nm.

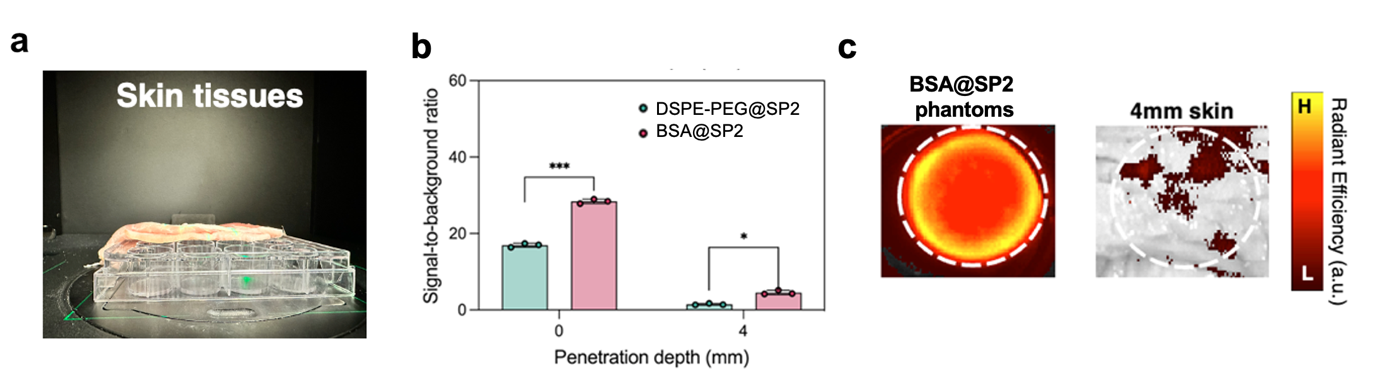

**Figure S19. Optical penetration of BSA@SP2 tumor-mimicking phantoms through porcine skin.** (a) White light image of porcine skin slices above tumor mimicking phantoms in a 12-well plate. (b) Fluorescence signal-to-background ratio and (c) IVIS images of tissue-covered tumor mimicking phantoms (n=3). IVIS images were collected at λ_ex_: 675 nm and λ_em_: 840 nm.
